# In Silico Characterization of a Key Cell Wall Enzyme for targeting Methicillin-Resistant *Staphylococcus aureus* using Bioactive compounds derived from *Haematococcus pluvialis*

**DOI:** 10.1101/2025.09.04.674297

**Authors:** S. Shakthi Bhavanee, T. P. Rajarajan, P. Hari Babu

## Abstract

Methicillin-resistant *Staphylococcus aureus* (MRSA) is a critical global problem in case of infection due to its multidrug resistance and adaptive nature. Polyisoprenyl-teichoic acid--peptidoglycan teichoic acid transferase (PTA), an essential enzyme for biosynthesis of teichoic acid is present in almost all of the gram-positive species, especially S. aureus. A biocompatible gel-patch formulated using acetone extract of microalgae *Haematococcus pluvialis* was developed and evaluated for its favourable physicochemical properties such as moisture retention and thickness. Antioxidant activity of the patch and extract was assessed via DPPH assay and results showed a dose-dependent radical scavenging potential and an IC₅₀ of 24.23 µg/mL, while antibacterial assay against *S. aureus* showed decent inhibition zones for the extract (5 - 7 mm) and the gel-patch (1 - 3.5 mm). In addition to invitro assays, in-silico analysis was carried out for predicting the effective usage of *H. pluvialis* against S. aureus strains. Conserved domain analysis identified the presence of a Cps2A domain of the LytR-CpsA-Psr (LCP) family in PTA. Physicochemical profiling revealed a hydrophilic, stable protein nature having high aliphatic index (85.07), low GRAVY score (–0.679), and a structured catalytic core with flanked disordered terminal regions. GC-MS analysis showed the presence of 15 compounds in the algal extract and ADMET analysis identified compounds with suitable drug-likeness, skin permeability and low systemic toxicity. The diversified residue-specific interactions for control drugs and the algal ligands in the conserved Cps2A domain favour a dual binding and synergistic therapeutic approach to target MRSA using the *H. pluvialis*-based gel-patch.

## 1. Introduction

Gangrene, a pathological condition characterized by ischemic tissue death is often resulting from inadequate blood supply, infection, or a combination of both (Al Wahbi, A., 2018). Fournier’s gangrene is a rare type of gangrene which is rapidly progressive and fatally necrotizing infection, affecting the perineal and genital regions. The illness primarily affects men and people who have conditions like immunodeficiency, diabetes mellitus, alcoholism and cancer. Moreover, a study analyzing 1,641 male and 39 female FG cases reported diabetes and obesity as significant risk factors for FG, whereas hypertension and substance use showed no significant association (Sorensen. M. D, & Krieger. J. N., 2016). A recently accomplished work focusing on microbial distribution and susceptibility of 150 patients had 45 patients with poly microbial infections, with *S. aureus* emerging as the predominant pathogen, owing to 76% of the total isolates, followed by *P. aeruginosa* and *E. coli* (Liu, W, et al., 2023). Diabetes significantly increases the risk of infections caused by Staphylococcus aureus across multiple organ systems and also, S. aureus emerged as the key pathogen in mucous membrane infections, contributing to folliculitis, furunculosis, and cellulitis (Muller LM, et al., 2005; Atreja A, Kalra S., 2015).

Even though numerous drugs are available for treating bacterial infections, MRSA (Methicillin Resistant Staphylococcus aureus) strains often neglect such drugs due to them harbouring genes that give resistance to most β-lactam antibiotics including methicillin. With combination to this resistance, S. aureus possesses virulence factors which increase the severity of infections (Lee. A. S, et al., 2018). Conventional wet-to-dry dressings and Negative Pressure Wound Therapy (NPWT) has emerged to be viable method to accelerate wound healing, decrease bacterial burden and promotes formation of granulation tissue. However, challenges persist in cases having extensive perianal involvement, where achieving effective negative pressure therapy can be difficult (Ozkan, O. F, et al., 2016).

LytR-CpsA-Psr (LCP) are a group of enzymes which catalyze the binding of undecaprenollinked intermediates to peptidoglycan, thereby anchoring the secondary cell wall polymers such as wall teichoic acid (WTA) in Staphylococcus aureus and capsular polysaccharides (CPS) in streptococci. LCP enzymes are vital for the anchoring of bacterial capsules and teichoic acids, which are possibilities to target in antimicrobial approaches. (Chan Y. G, et al., 2014). Scientists tabulated a complete list of LCP proteins and surveyed their phylogenetic distribution, and found its presence in all Gram-positive bacteria with the exception of Mollicutes and one species of Clostridiales, while most of the Gramnegative bacteria did not have these proteins. Experimental verification with Staphylococcus aureus MsrR validated the inferred membrane topology and extracellular location of the LCP domain, indicating that the LytR-CpsA-Psr domain is specific to bacteria and found in various subgroups, which could be responsible for the functional diversity seen in various bacterial species (Hübscher J, et al., 2008). S. aureus contains three lcp genes, and their deletion (Δlcp) causes WTA release and growth defects (Chan Y. G, et al., 2014). The enzyme polyisoprenyl-teichoic acid--peptidoglycan teichoic acid transferase is essential for Staphylococcus aureus wall teichoic acid (WTA) biosynthesis, facilitating the final transfer of teichoic acid precursors from a lipid carrier to peptidoglycan, ensuring cell wall integrity, morphology, and growth (Keinhörster D, et al., 2019). Inhibiting this enzyme presents a promising antibacterial strategy, as WTA disruption impairs S. aureus division, reduces virulence by hindering host adhesion and immune evasion, and increases susceptibility to β-lactam antibiotics (Keinhörster D, et al., 2019; Winstel V, et al., 2014).

Microalgae are widely utilized in the biomedical and pharmaceutical sectors due to their diverse bioactive compounds (Kumar, T. S, et al., 2023). Haematococcus pluvialis is a unicellular, spherical and biflagellate microalga, which, depending on its life cycle, morphology, and physiology, can exist in either vegetative green cells or red cysts. The green cells are associated with the role of biomass accumulation, are motile due to their two flagella, while the red cysts accumulate astaxanthin under stress conditions such as nutrient deprivation or exposure to intense light (Butler, T, et al., 2018). Astaxanthin, a significant carotenoid which gets accumulated exclusively during the red phase, owing to 80%–99% of the total carotenoids (Dragos, N, et al., 2010). Astaxanthin exhibits high antioxidant activity by scavenging free radicals, stabilizes biological membranes and prevents oxidative stress damage in cells. Despite its potential, astaxanthin has limitations in bioavailability and stability (Kumar, T. S, et al., 2023). Moreover, all research predominantly emphasizes the red stage due to its high astaxanthin content, often overshadowing the therapeutic potential of the green stage. An investigation demonstrated the antihypercholesterolemic activity of astaxanthin extracted from green-stage H. pluvialis, suggesting its potential in managing cholesterol levels and exists unexplored. Additionally, Lutein is the dominant carotenoid in green cells, accounting for 70%–80%, followed by β-carotene (16.7% d.w.), violaxanthin (12.5% d.w.), neoxanthin (8.3% d.w.) and chlorophyll at 1.5%–2% d.w, and these compounds are either absent or present in minimal amounts in red-phase cells (Shah, M.M.R, et al., 2016). *H. pluvialis* is also rich in proteins, essential vitamins and minerals like calcium, magnesium, iron, and zinc (Kumar, T. S, et al., 2023), and the rich lutein and β-carotene content in green stage may offer antioxidant benefits pertinent to eye health and immune function (Ashaq Hussain Rather, Rekha Rao., 2020). Despite these findings, comprehensive research on the therapeutic applications of greenstage *H. pluvialis* remains limited, indicating a need for further exploration to fully elucidate its health benefits. In addition to given the lack of in-silico validation for polyisoprenyl-teichoic acid-peptidoglycan teichoic acid transferase in S. aureus, this work focuses on modelling the 3D structure of the protein and evaluate the anti-bacterial effects of Dichloromethane extract from *H. pluvialis* in it green cell stage using molecular docking and ADMET characterization.

## 2. Materials and Methods

Distilled water, Type A gelatin (porcine origin), polyethylene glycol (PEG-400), acetone, nutrient agar, DPPH (2,2-diphenyl-1-picrylhydrazyl), acetyl salicylic acid, petri dish, test tubes, test tube stands, beakers, Magnetic heating stirrer, UV-Vis spectrophotometer, OHM media, centrifuge,

### 2.1. Sample Collection and Preparation

Fresh *Hematococcus pluvialis* were purchased from local markets in Chennai and were cultivated for a period of 7 days to reach its peak green cell stage in OHM media prepared by standard method (imamoglu. E, et al., 2008). The culture was centrifuged for 5mins at 10000 rpm and the biomass was subjected to cold maceration (4 °C) with Acetone as the solvent for 24hrs. The solution is centrifuged for 10 mins at 5000 rpm and the extract was collected in a brown glass bottle for storage in sterile conditions (Miazek. K, et al., 2017).

### 2.2. Gel Formulation

The sprayable gel formulation from (Islam. M. S, et al., 2021) was slightly modified to be used as an instant gel patch by replacing the usage of diethyl ether with *H. pluvialis* acetone extract. The homogenous Gelatin-PEG solution (after stirring) is allowed to cool and algal extract is added in increasing concentrations and stirred continuously. It is then poured in petri dishes approximately 5mm thickness and allowed to dry at 37 °C for 30 – 120 seconds. The sheets were stored in foils and 4 °C for long-term storage and direct wound application.

### 2.3. Physical Characterization of Gel

*H. pluvialis* extract-loaded gel patch was assessed to evaluate its suitability for wound healing applications. The gel solidifying time, thickness of the gel post-drying, flexibility and folding through manual measurement were recorded. Additionally, endurance and moisture retention capacity of the patch were studied to maintain optimal wound environment.

### 2.4. Evaluation of Anti-Oxidant Activity - DPPH Assay

*H. pluvialis* acetone extract’s antioxidant capacity was assessed by measuring the potential to scavenge DPPH radicals. A solution containing 0.1 mM DPPH in methanol was added to 1 ml of varying concentrations (25, 50, 75, 100, 125 μg/ml) of algal extract. Control was prepared by mixing 1 ml solution of methanol, DPPH and acetone, and all solutions were left undisturbed in the dark for 30 mins. The percentage of inhibition of radicals was observed by measuring absorbance at 517 nm. The % radical scavenging activity is calculated using the formula:

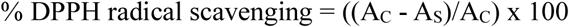

Where, A_C_ – Absorbance of control and A_S_ – Absorbance of sample at 517nm.

### 2.5. Evaluation of Antibacterial Activity

*H. pluvialis* extract and gel patch both were tested for their anti-bacterial potential against *Staphylococcus aureus* strain with acetyl salicylic acid as the control. Well diffusion method was carried out by preparing agar plates with 6 wells and 5 wells (6mm in diameter) for evaluation of 4 different concentrations (100 – 400 μg/ml) of algal extract and gel patch respectively. Control drug, acetyl salicylic acid (25 µg/ml) was used as the positive control and acetone/gel patch without extract, was used as the negative control. The plates were incubated at 37°C for 24 hrs and the formed inhibition zones were observed and measured.

### 2.6. Gas chromatography–Mass Spectrometry

An Agilent 6890N JEOL GC Mate II system with an HP-5 column (30 m × 0.25 mm, 0.25 μm film thickness) was used for the GC-MS with helium as carrier gas at a flow rate of 1 mL/min. The column temperature was set at 200°C and conditions for mass spectrometry included a mass scan range of 50– 600 m/z, an ionization voltage of 70 eV, and source and interface temperatures of 250°C. The collected spectra were compared with entries in the NIST collection in order to identify the compound (Yücel. T. B, & Yaylı. N., 2018)

### 2.7. ADMET characterization

The GC-MS compounds were searched in pubchem compound database to acquire their smile formats and sdf structures. The SwissADME server was utilized for predicting the pharmacokinetic properties of bioactive compounds, while Molinspiration was used to evaluate their drug likeness (Seyed. M. A, et al., 2022). Protox-III server was used to predict toxicological endpoints and toxicity degree (LD50, mg Kg^-1^) of the bioactive compounds (Banerjee. P, et al., 2024), with additional insights using GUSAR (General Unrestricted Structure–Activity Relationships) to predict the acute toxicity data and subcutaneous LD₅₀ values classified according to the OECD chemical toxicity classification system (Savjani. J, et al., 2022). Further evaluation of dermal toxicity, skin permeation, irritation, sensitization, and environmental biodegradability using VEGA QSAR platform to provide a comprehensive in-silico validation (Hong-sheng. M. A, et al., 2022).

### 2.8. Evaluation of Physico-chemical properties

The amino acid sequence of Polyisoprenyl-Teichoic Acid (Accession number: WP_412521717.1) was acquired using NCBI database, and it was utilized to investigate the protein’s atomic composition, relative molar mass, isoelectric point (pI), and other physiochemical properties were calculated utilizing the ExPASy ProtParam software (Raen. R, et al., 2024).

### 2.9. Protein Structure Prediction and Comparative Analysis

The secondary structure characteristics were analyzed using SOPMA program with default parameters (Raen. R, et al., 2024) and the degree of disorderness and sequence analysis was performed using PSIPRED (Buchan. D. W, et al., 2024). For 3D structure predition, the amino acid sequence of PTA was subjected to BLAST against the Protein Data Bank (PDB) structure to get structurally resolved homologs (Bitencourt-Ferreira. G, et al., 2019) and their corresponding PDB 3D structures were retrieved from the RCSB PDB database to be used as templates. The initial models were refined through loop modeling and optimization techniques to enhance structural accuracy. In addition, SwissModel (Schwede. T, et al., 2003), AlphaFold (Roney. J. P, & Ovchinnikov. S., 2022), Phyre2 (Kelley. L. A, et al., 2015) and Rosetta (Wang. W, et al., 2022) were utilized 3D structural models to ensure the best prediction accuracy and comparative selection.

### 2.10. In-silico Analysis

#### 2.10.1. Ligand and protein preparation

The generated models were refined using Galaxy Refine and the enhanced protein models were evaluated to select the best model, which is used for subsequent molecular docking studies (Heo. L, et al., 2013). The final model in its PDB format was checked with any abnormalities/missing residues, were corrected, and prepared for docking using Dockprep (Amber20 Forcefield) in ChimeraX (Kumari. M., 2023). On the other hand, the ligand structures in SDF file format were performed with energy minimization using Open Babel and these structures were converted to pdbqt format for docking (Rekha. U. V, et al., 2022).

#### 2.10.2. Docking Analysis and Visualization of Interactions

Molecular docking simulations were conducted using PyRx software and the results were assessed using the calculated binding affinities (kcal/mol) and ligand-protein interactions (Rekha. U. V, et al., 2022). Commercial drugs Chloramphenicol (PubChem Id: 5959) and Gentamycin (Pubchem Id: 3467) were used for blind docking. Ligand conformation with the lowest binding energy and the smallest RMSD values were selected for displaying the binding sites and interaction profile using Discovery Studio BIOVIA (Akash. S, et al., 2023). To determine potential ligand-binding sites, Computed Atlas of Surface Topography of Proteins (CASTp) server was employed to analyze the cavities/pockets present in the protein (Chandran. K., 2022). Both Blind docking and CastP data were analysed to predict the active sites of the protein. Based on this, the functional active site regions were selected to enhance docking accuracy and computational efficiency.

## 3. Results and Discussion

### 3.1. Algal Gel-Patch Compatibility

The substitution of diethyl ether (DEE) with acetone extract created a stable and solid patch which gelled instantly upon casting. Gelation occurred within 30–60 seconds after pouring into petri plate and additionally facilitated quick evaporation of acetone, leaving the bioactive algal compounds etched in the patch. This new formulation being slow in solidification compared to the rapid 3 second gelation of the DEE spray, addresses previous concerns related to toxicity and volatility of DEE for the spray. Uniform distribution of the extract throughout the gel matrix ensured no disturbance in the gel matrix integrity.

The *gel-*patch showed a decrease in thickness from 5 mm on the time of casting to 3.5 mm after letting it dry for 6 hrs. Both algal gel-patch and the DEE spray had gelatin’s moisture-sealing properties. However, the patch had greater control over hydration and vapor permeability with the moisture content reduced by around 24% during the drying period (Figure 1.A), with evaporation of acetone, leaving the bioactive compounds in the gel. The gel-patch can also be layered or trimmed making it customizable based on wound coverage and managing chronic or diabetic wounds.

**Fig.1.**
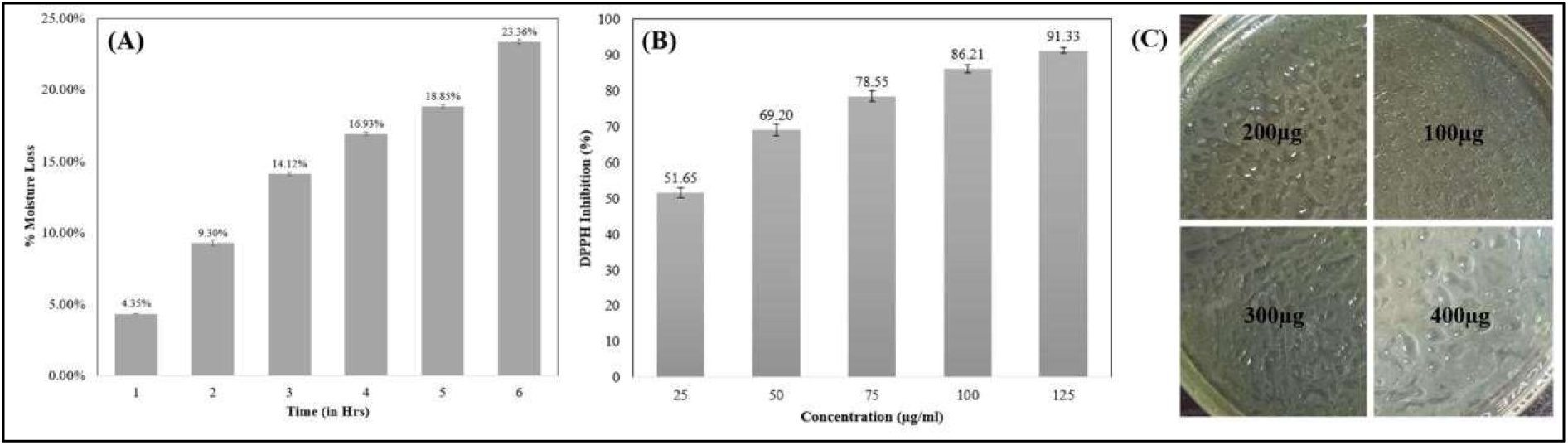
Evaluation of *H. pluvialis* Acetone Extract and Gel-Patch: (A) Loss in Moisture Assessed Over a Period After Gel Casting, (B) Antioxidant Activity of *H. pluvialis* Acetone Extract via DPPH assay, (C) Visualization of Casted Gel at 4 Concentrations.

Additionally, the patch was observed to be fresh (No rotting or dehydration) for a period of 6 months when stored in sealed, light-proof pouches at 4 °C. The DEE spray required careful packaging and its high volatility and flammability adds concerns to storage and transportation. With enhanced shelf-life and minimal/absence of solvent residues, the gel-patch offers improved safety margins, reduces risks of skin irritation, provides cost-effective and feasible application.

### 3.2. Anti-Oxidant Activity using DPPH Assay

DPPH free radical scavenging assay showed the algal acetone extract to possess a dosedependent antioxidant activity (Figure 1.B). At 25 µg/mL, the extract exhibited 51.65% inhibition activity and progressively rose to 69.20%, 78.55%, and 86.21% at 50, 75 and 100 µg/mL respectively, with maximum inhibition of 91.33% observed at 125 µg/ml. With an estimated IC₅₀ value of 24.23 µg/mL, *H. pluvialis* demonstrated such potent antioxidant potential due to the presence of polyphenols, carotenoids, etc. The concentration dependent antioxidant activity further makes it suitable for reducing oxidative stress and therapeutic applications using optimized formulations.

### 3.3. Antibacterial Activity

The antimicrobial potential of the *H. pluvialis* extract was quantitatively assessed by measuring the zones of inhibition produced against S. aureus (Figure 1.D). The extract showed a dosage wise increase in antibacterial activity with inhibition zones ranging from 5.13 ± 0.06 mm (100 µg/ml) to 7.40 ± 0.00 mm (400 µg/ml). However, the standard positive control had 9.97 ± 0.06 mm zone of inhibition, indicating a moderate level difference for the potent antibacterial efficacy of the algal extracts.

The gel patch coated with the extract showed similar consistency but reduced inhibition zones ranging from 1.03 ± 0.06 mm for 100 µg/ml to 3.47 ± 0.06 mm for 400 µg/ml concentration. This reduction in activity is either reduction in absorption of the compounds or due to slower diffusion of the bioactive compounds from the gel matrix. The unloaded gel patch showed minimal inhibition (0.07 ± 0.06 mm) and solvent control showed a moderate inhibition (4.83 ± 0.06 mm), validating *H. pluvialis* for its antimicrobial action. These results correlate with the results of various solvent extracts of red cell stage H. pluvialis, where in various solvents, where chloroform and ethyl acetate extracts showed the strongest antibacterial activity, particularly against B. subtilis (17.32 mm) and Escherichia coli (15.87 mm), while acetone and methanol extracts exhibited moderate effects, notably against Listeria monocytogenes (13.96 mm, 13.18 mm), suggesting *H. pluvialis* as potential bacteriostatic agents (Rao, A, et al., 2010). The DEE spray exhibited antibacterial activity with 6–10 mm zones of inhibition sue to DEE’s antimicrobial properties. However, due to the reduced toxicity, zero skin irritation and decent anti-bacterial activity of *H. pluvialis* extract loaded gel-patch makes it feasible and biocompatible for wound care and controlled bioactive compound release.

**Table.1.** Antibacterial Activity Assessment of *H. pluvialis* Acetone Extract and Extract Loaded Gel-Patch Against *S. aureus*.

| Sample (or)<br>Concentration ($\mu\text{g}/\text{well}$) | Extract<br>Zone of Inhibition (mm) | Gel Patch Zone of<br>Inhibition (mm) |
| --- | --- | --- |
| Acetone (99%)/Unloaded<br>Gel Patch | $4.83 \pm 0.06$ | $0.07 \pm 0.06$ |
| 100 | $5.13 \pm 0.06$ | $1.03 \pm 0.06$ |
| 200 | $5.83 \pm 0.12$ | $1.97 \pm 0.12$ |
| 300 | $6.67 \pm 0.06$ | $2.80 \pm 0.00$ |
| 400 | $7.40 \pm 0.00$ | $3.47 \pm 0.06$ |
| Standard | $9.97 \pm 0.06$ | — |

### 3.4. Computational Analysis of the Protein Sequence

The target protein, WP_412521717.1, from *Staphylococcus aureus*, is a full-length protein with 408 amino acids and a calculated molecular weight of ∼45.9 kDa. Annotated as a polyisoprenyl-teichoic acid--peptidoglycan teichoic acid transferase, CDD analysis revealed the presence of a prominent conserved region spanning residues 62 to 310, matching the Cps2A domain (COG1316), which is a part of the LytR-CpsA-Psr (LCP) family, known for its involvement in anionic cell wall polymer biosynthesis and attachment of teichoic acids to peptidoglycan layers (Chan Y. G, et al., 2014). Further analysis showed strong similarity to the Lytr-CpsA-psr superfamily, which supports a bifunctional role in enzymatic transferase activity and transcriptional regulation within bacterial systems (Keinhörster D, et al., 2019; Winstel V, et al., 2014). Specific domain matches such as PRK09379 and TIGR00350 reinforce the dual functional nature, suggesting both structural and regulatory functions in bacterial envelope formation. The unique presence of LCP domains in bacterial systems, and their absence in humans, provides a selective therapeutic window. Inhibiting such transferases could disrupt cell wall integrity without affecting host cells, making WP_412521717.1 a viable candidate for novel antimicrobial drug development (Winstel V, et al., 2014; Keinhörster D, et al., 2019).

**Fig.2.**
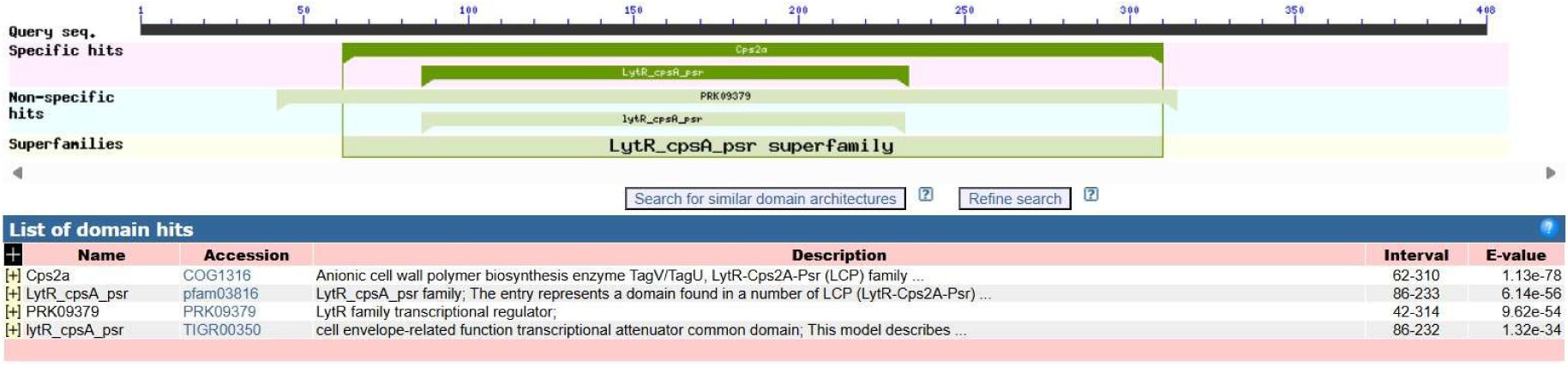
Conserved Domain Analysis of Target Protein (PTA) Using NCBI CD-Search.

#### 3.4.1. Physicochemical Characterization

The physicochemical characterization of the target protein revealed several important insights corresponding to its nature and functions (Table. 2). The theoretical pI was calculated as 6.16, suggesting that the protein would exhibit a net negative charge under physiological pH conditions (∼7.4). The number of negatively charged residues (52) slightly exceeded the positively charged residues (48), which aligns with the slightly acidic pI. Protein stability was evaluated through the instability index (II), which yielded a value of 35.25 (<40), indicating it to be stable under in vitro conditions.

**Table.2.** Physicochemical Properties of the Target Protein (PTA) Predicted Using ProtParam.

| Category | Parameter | Value |
| --- | --- | --- |
| **Basic Properties** | Number of amino acids | 408 |
|  | Molecular weight | 46,041.65 Da |
|  | Theoretical pI | 6.16 |
|  | Total number of atoms | 6,448 |
|  | Negatively charged residues (Asp + Glu) | 52 |
|  | Positively charged residues (Arg + Lys) | 48 |
| **Extinction Coefficient (280 nm)** | M<sup>-1</sup> cm<sup>-1</sup> | 25,330 |
|  | Absorbance (0.1% = 1 g/L) | 0.550 |
|  | Note | No Trp residues; >10% error |
| **Estimated Half-life** | Mammalian reticulocytes (in vitro) | 30 hours |
|  | Yeast (in vivo) | >20 hours |
|  | *E. coli* (in vivo) | >10 hours |
| **Stability & Hydrophobicity** | Instability index (II) | 35.25 (Stable) |
|  | Aliphatic index | 85.07 |
|  | GRAVY (Hydropathicity) | -0.679 (Hydrophilic) |

The aliphatic index was relatively high at 85.07, suggesting that the protein may retain structural integrity across a broad temperature range, an important feature for industrial or environmental applications. The GRAVY score was -0.679, indicating a predominantly hydrophilic nature (Saikia. A, et al., 2021). Lastly, the estimated half-life across systems like mammalian reticulocytes (30 hours), yeast (>20 hours), and *E. coli* (>10 hours) underscores the protein’s relative stability and persistence across different expression hosts.

The amino acid composition was evaluated and it showed an abundance of aspartic acid (10.0%), leucine (9.1%), and lysine (8.1%), followed by serine (7.1%), glutamic acid (6.6%) and alanine/valine (5.9% each). These residues could contribute significantly to the protein’s overall charge, solubility, and structural conformation. Furthermore, the high levels of charged residues (Asp, Glu, Lys) correlate with the hydrophilic GRAVY score (Saikia. A, et al., 2021) and further support the coil-rich and flexible nature of the protein.

**Fig.3.**
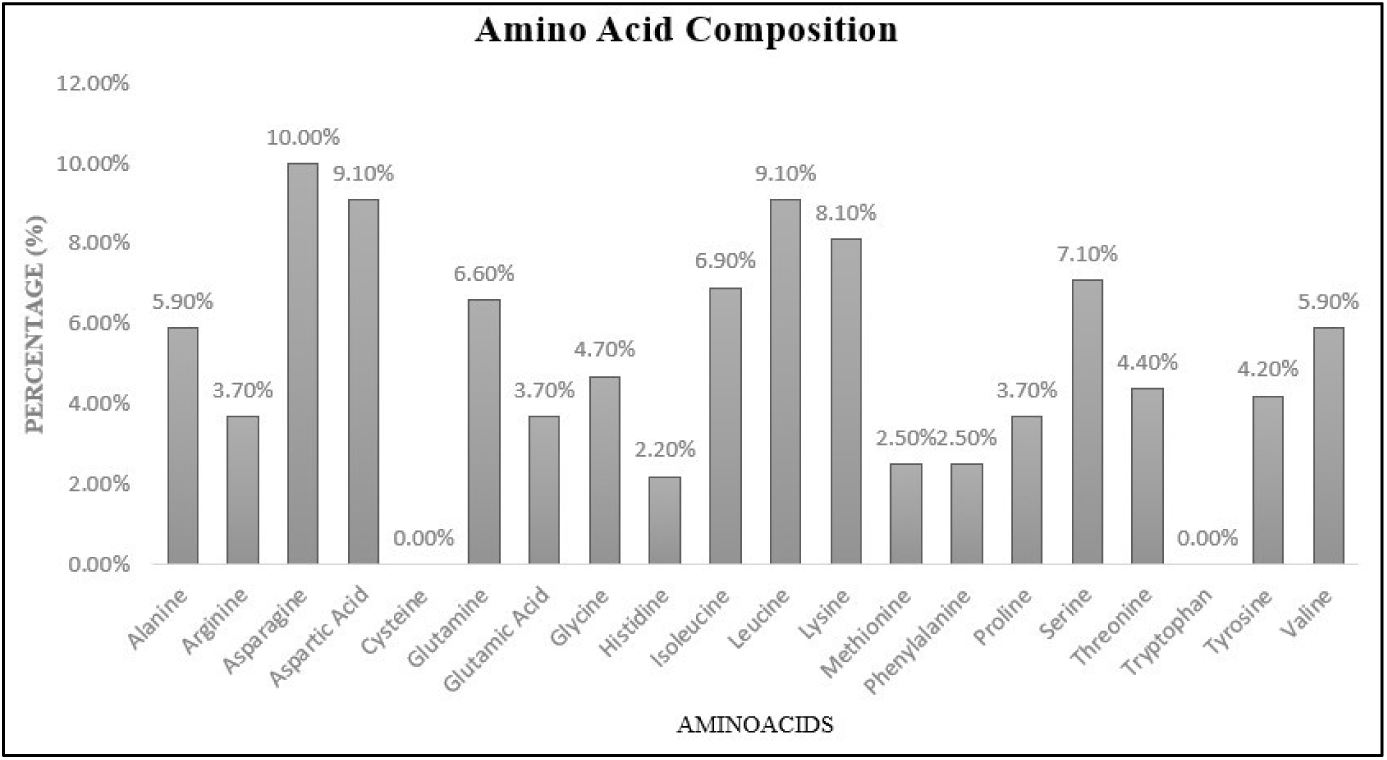
Amino Acid Composition of the Target Protein determined using Protparam.

#### 3.4.2. Secondary Structure Prediction and Analysis

Secondary structure prediction using SOPMA indicated that the protein was majorly composed of random coils and α-helices, with only a small portion forming β-strands. The structural composition was predicted as α-Helix: 130 residues (31.86%), Extended Strand (β-sheet): 49 residues (12.01%) and Random Coil: 229 residues (56.13%) with no 3_10_ or π helices, β-bridges, or β-turns. This suggests that the protein adopts a flexible structure with a considerable portion remaining disordered or unstructured, especially at the termini. However, the structured core within residues 62–310 (Cps2A domain), is indicative of active domain architecture suitable for enzymatic catalysis. Moreover, the predicted Nterminal helices might contribute to membrane association. Hence, the secondary structure of the target protein was further analyzed using the PSIPRED tool, providing a detailed and high-confidence prediction.

From the PSIPRED output, it was observed that α-helices were prominently distributed in residues between 100 and 300, i.e., the central regions of the protein, while β-strands appeared to be more frequent between residues ∼50–250. Random coils were now confirmed to dominate the N- and C-terminal regions and in between the helices and strands, underscoring that the protein might be flexible. The confidence score bar in blue further indicated that the central domain has a high prediction reliability.

**Fig.4.**
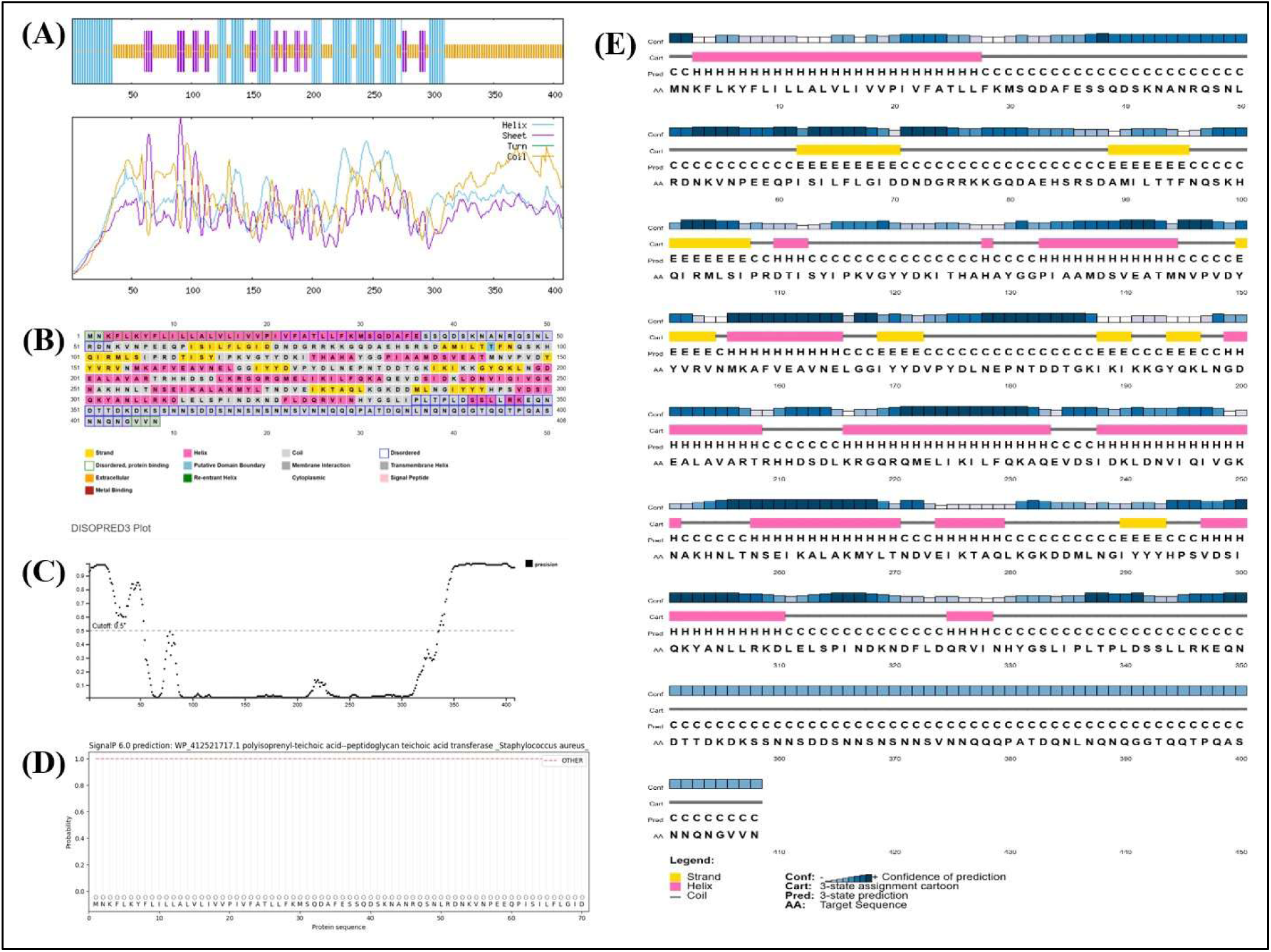
Comprehensive Structural Analysis of PTA using (A) SOPMA-Based Secondary Structure Prediction, (B) Amino Acid Profile via PSIPRED, (C) Intrinsic Disorder Prediction, (D) Signal Peptide prediction in the protein using SignalP, and (E) Secondary Structure Confidence Assessment Using DMPFold 2.0.

DISOPRED3 plot further revealed the N-terminal region (residues ∼1–90) and C-terminal region (residues ∼320–408) as the two major disordered regions. Further strengthening this hypothesis, a strong N-terminal signal peptide (MNKFLKYFLILLALVLIVVPIVF) was identified, which is highly hydrophobic and typically directs the protein towards secretion pathways or membrane insertion. These overlapping results from SOPMA, PSIPRED and DISOPRED confirm the presence of a bipartite nature in the protein, where Ordered Core (residues ∼100–300) is likely responsible for the protein’s main structural or catalytic functions, while Flexible Termini (residues ∼1–90 and 320–408) are involved in molecular recognition, interactions or regulatory mechanisms (Haliloglu. T, & Bahar. I., 2015).

### 3.5. Comparative Modeling, Refinement, and Validation of the Target Protein

Five models were generated using the templates T1-T5 with the Modeller software. Based on the weighted pair-group average clustering, the model T3A (score: 1.0000) had the closest structural similarity, followed by T2A (score: 66.2500) and T4A (score: 70.5000), which were more distant. Each model was evaluated using DOPE and GA341 scores to ensure the highest possible quality for downstream analysis. The GA341 scores for the selected models ranged between 0.99998 and 1.00000, indicating near-perfect accuracy and stability in the models.

The DOPE score for 4^th^ model of template 4 (T4A) was -23642.64844, which was the most favorable among the models, suggesting a highly stable structure. The GA341 score for this model was 1.00000, indicating it was highly reliable (Bitencourt-Ferreira. G, et al., 2019). This model was further analyzed in Chimera and was confirmed to have key regions such as the binding sites and active sites, thereby showing good compatibility with that of the target protein. For Swissmodel, the final model was selected on the basis of QMEAN, GMQE, MolProbity and Ramachandran plot analyses (Saikia. A, et al., 2021). Model 2 was selected as the best candidate as it exhibited the highest percentage of Ramachandran favored residues (94.72%), least Ramachandran outliers (0.38%) and rotamer outliers (0.43%).

**Fig.5.**
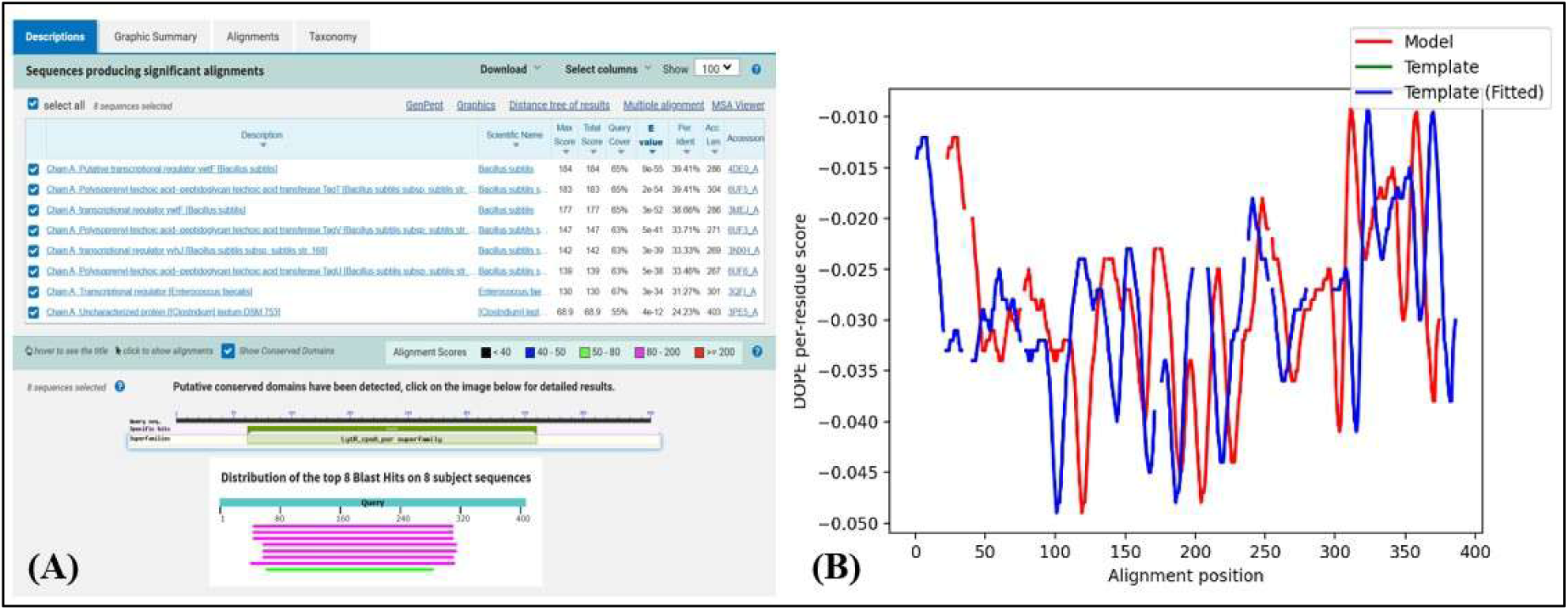
(A) BLASTp Analysis for PTA across Related Bacterial Sequences and (B) Corresponding DOPE Energy Profile of the Modelled Protein.

A comparative evaluation of the protein models generated from different prediction tools were performed after refinement and the resulting models reveal significant differences in structural accuracy and stereochemical quality, which are summarized in Table 3. Rosetta protein model emerged as the best-performing model overall, as it covered the entire protein sequence and the refined model had a GDT-HA of 0.9798 and the lowest RMSD of 0.328, indicating the highest atomic-level accuracy. It also maintained a strong stereochemical profile, with a MolProbity score of 1.449, Clash score of 8.3, and 98.3% of residues in the Ramachandran favored region. However, AlphaFold presented the most refined stereochemistry, achieving the lowest MolProbity score (1.348) and Clash score (6.3), with 0% poor rotamers. Despite its slightly higher RMSD (0.420), the superior structural refinement made it the most suitable candidate for molecular docking studies and interaction analysis. Alphafold structure was chosen for further docking studies and was prepared through standardization of charges and atom types using the Amber 20 force field, ensuring compatibility with energy minimization and docking protocols.

**Fig.6.**
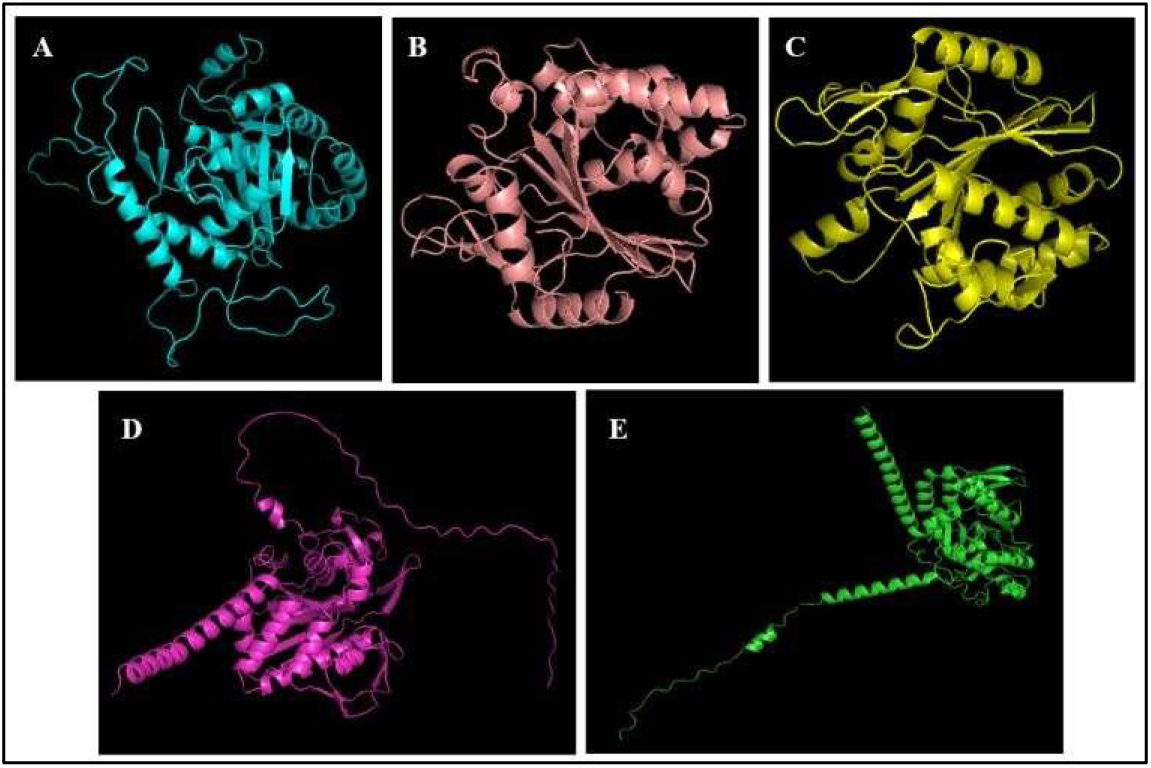
Visualization of the best 3D Protein Structure models generated by (A) Modeller, (B) SWISS-MODEL, (C) Phyre2, (D) AlphaFold and (E) Rosetta.

**Table.3.** Comparative Evaluation of Refined Protein Models based on Structural and Stereochemical Properties.

| Model | GDT-HA | RMSD | MolProbity | Clash score | Poor rotamers | Rama favored |
| --- | --- | --- | --- | --- | --- | --- |
| Advanced HM | 0.9534 | 0.424 | 1.585 | 8.4 | 0.3 | 97.3 |
| Rosetta | 0.9798 | 0.328 | 1.449 | 8.3 | 0.5 | 98.3 |
| AlphaFold | 0.9694 | 0.420 | 1.348 | 6.3 | 0.0 | 98.0 |
| Phyre | 0.9647 | 0.402 | 1.627 | 13.1 | 0.4 | 98.1 |
| Swissmodel | 0.9663 | 0.393 | 1.495 | 9.3 | 0.4 | 98.9 |

### 3.6. ADMET Analysis

The bioactive profile of *H. pluvialis* extracted using Acetone revealed 15 distinct bioactive compounds through GC-MS analysis. Among the detected metabolites, palmitic acid (n-hexadecanoic acid), oleic acid and linoleic acid are well known for their antimicrobial activity by disrupting microbial integrity and anti-inflammatory properties. Apart from this, several bioactive esters and heterocycles were also detected, providing a hint for the therapeutic relevance of *H. pluvialis* in treating chronic wounds. Further evaluation of ADME properties and Toxicity of the ligands area evaluated.

**Table.4.**
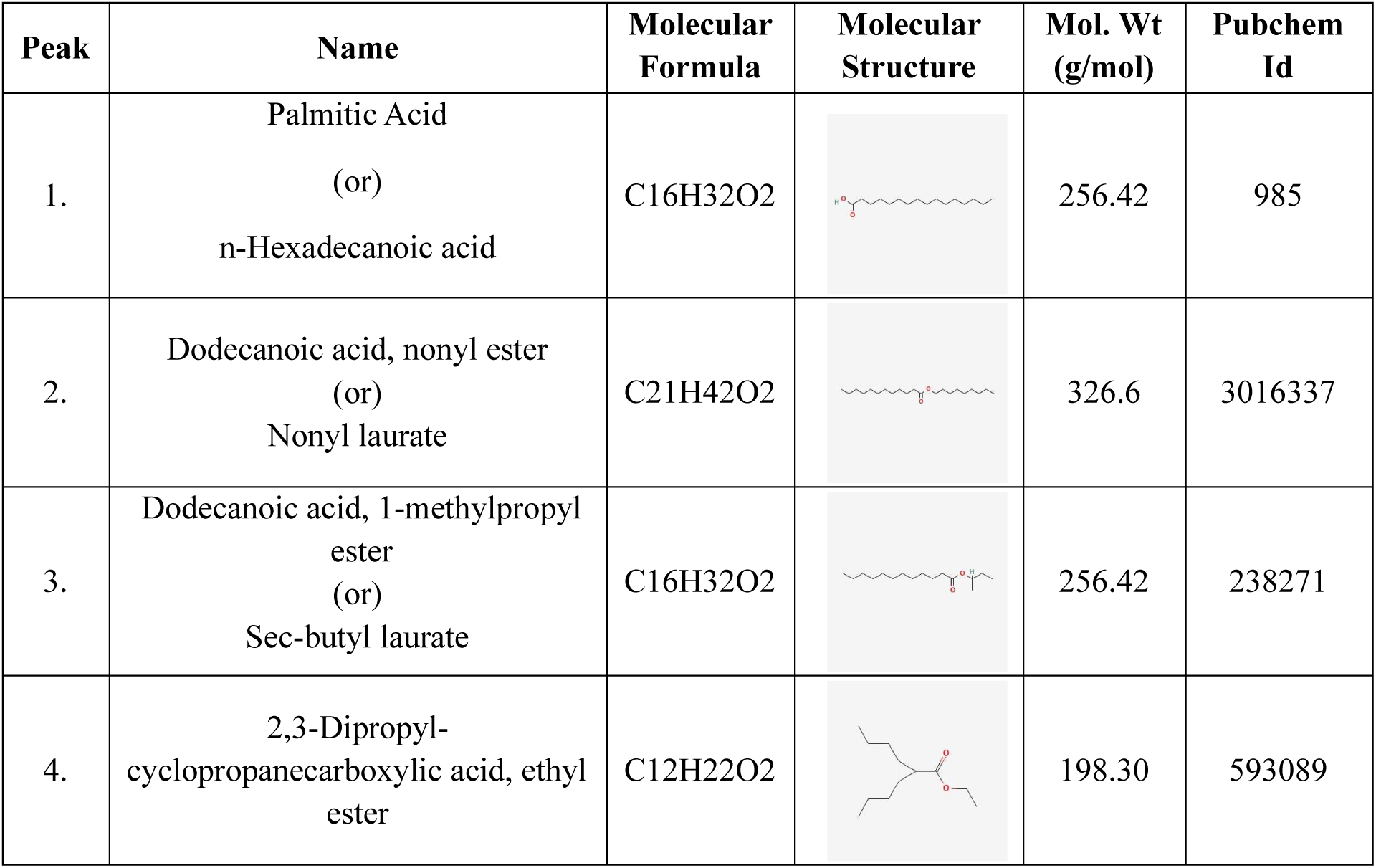
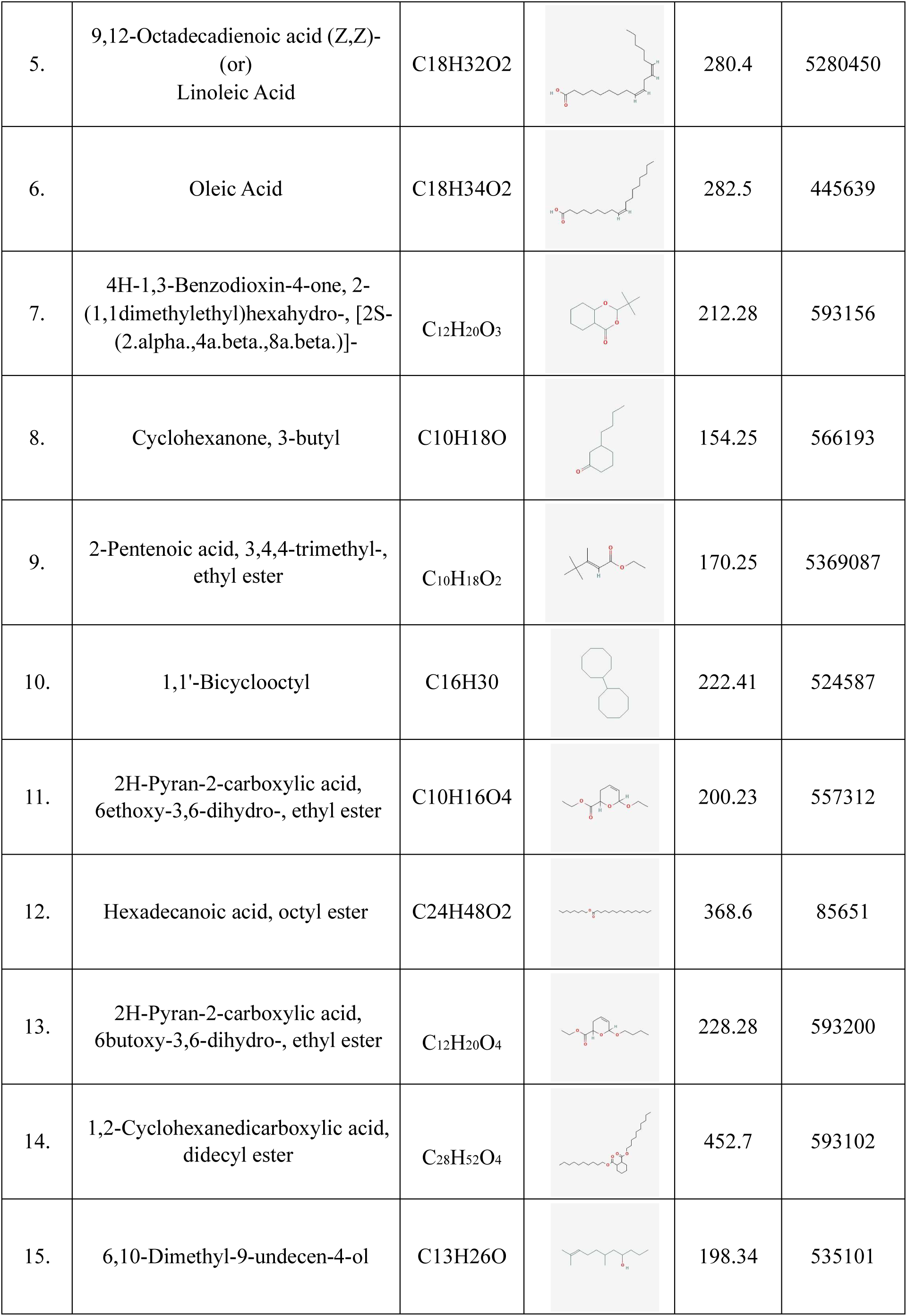
Bioactive Compounds Identified by GC-MS from Acetone Extracts of Green-stage *H. pluvialis*.

| Peak | Name | Molecular Formula | Molecular Structure | Mol. Wt (g/mol) | Pubchem Id |
| --- | --- | --- | --- | --- | --- |
| 1. | Palmitic Acid<br>(or)<br>n-Hexadecanoic acid | C16H32O2 |  | 256.42 | 985 |
| 2. | Dodecanoic acid, nonyl ester<br>(or)<br>Nonyl laurate | C21H42O2 |  | 326.6 | 3016337 |
| 3. | Dodecanoic acid, 1-methylpropyl ester<br>(or)<br>Sec-butyl laurate | C16H32O2 |  | 256.42 | 238271 |
| 4. | 2,3-Dipropyl-cyclopropanecarboxylic acid, ethyl ester | C12H22O2 |  | 198.30 | 593089 |
| 5. | 9,12-Octadecadienoic acid (Z,Z)-<br>(or)<br>Linoleic Acid | C<sub>18</sub>H<sub>32</sub>O<sub>2</sub> |  | 280.4 | 5280450 |
| 6. | Oleic Acid | C<sub>18</sub>H<sub>34</sub>O<sub>2</sub> |  | 282.5 | 445639 |
| 7. | 4H-1,3-Benzodioxin-4-one, 2-(1,1dimethylethyl)hexahydro-, [2S-(2.alpha.,4a.beta.,8a.beta.)]- | C<sub>12</sub>H<sub>20</sub>O<sub>3</sub> |  | 212.28 | 593156 |
| 8. | Cyclohexanone, 3-butyl | C<sub>10</sub>H<sub>18</sub>O |  | 154.25 | 566193 |
| 9. | 2-Pentenoic acid, 3,4,4-trimethyl-, ethyl ester | C<sub>10</sub>H<sub>18</sub>O<sub>2</sub> |  | 170.25 | 5369087 |
| 10. | 1,1'-Bicyclooctyl | C<sub>16</sub>H<sub>30</sub> |  | 222.41 | 524587 |
| 11. | 2H-Pyran-2-carboxylic acid, 6ethoxy-3,6-dihydro-, ethyl ester | C<sub>10</sub>H<sub>16</sub>O<sub>4</sub> |  | 200.23 | 557312 |
| 12. | Hexadecanoic acid, octyl ester | C<sub>24</sub>H<sub>48</sub>O<sub>2</sub> |  | 368.6 | 85651 |
| 13. | 2H-Pyran-2-carboxylic acid, 6butoxy-3,6-dihydro-, ethyl ester | C<sub>12</sub>H<sub>20</sub>O<sub>4</sub> |  | 228.28 | 593200 |
| 14. | 1,2-Cyclohexanedicarboxylic acid, didecyl ester | C<sub>28</sub>H<sub>52</sub>O<sub>4</sub> |  | 452.7 | 593102 |
| 15. | 6,10-Dimethyl-9-undecen-4-ol | C<sub>13</sub>H<sub>26</sub>O |  | 198.34 | 535101 |

#### 3.6.1. Lipinski’s rule for drug likeness (Molinspiration)

The physicochemical properties including miLogP, Topological Polar Surface Area (TPSA), molecular weight (MW), hydrogen bond donors and acceptors, and other drug-likeness parameters were assessed to evaluate their suitability for oral/topical applications and potential antibacterial activity, particularly against *Staphylococcus aureus* (Seyed. M. A, et al., 2022; Rukthong. P, et al., 2020). Most compounds showed high lipophilicity, with hexadecanoic acid, octyl ester and 1,2Cyclohexanedicarboxylic acid, didecyl ester exhibiting the highest miLogP value of 9.34 and 9.42 respectively. Increased lipophilicity is beneficial for topical applications (skin penetration) but hinders oral bioavailability. On the other hand, 2H-pyran-based esters showed low miLogP values (<2.6), indicating better solubility in aqueous environment.

**Table.5.** Physicochemical Properties of Selected Ligands from Plant extract stored at ambient temperature.

| Compound Name | miLogP | TPSA | natoms | MW | nON | nOHNH | nviolations | nrothb | Volume |
| --- | --- | --- | --- | --- | --- | --- | --- | --- | --- |
| Palmitic Acid (n-Hexadecanoic acid) | 7.06 | 37.30 | 18 | 256.43 | 2 | 1 | 1 | 14 | 291.42 |
| Dodecanoic acid, nonyl ester (Nonyl laurate) | 8.84 | 26.30 | 23 | 326.56 | 2 | 0 | 1 | 19 | 376.16 |
| Dodecanoic acid, 1methylpropyl ester (Sec-butyl laurate) | 6.59 | 26.30 | 18 | 256.43 | 2 | 0 | 1 | 13 | 291.93 |
| 2,3-Dipropylcyclopropanecarboxylic acid, ethyl ester | 4.76 | 26.30 | 14 | 198.31 | 2 | 0 | 0 | 7 | 213.94 |
| Linoleic Acid (9,12-Octadecadienoic Acid) | 6.86 | 37.30 | 20 | 280.45 | 2 | 1 | 1 | 14 | 312.65 |
| Oleic Acid (9-Octadecenoic acid) | 7.58 | 37.30 | 20 | 282.47 | 2 | 1 | 1 | 15 | 318.84 |
| 4H-1,3-Benzodioxin-4-one, 2-(1,1dimethylethyl)hexahydro-, [2S(2α,4$\alpha\beta$,8$\alpha\beta$)] | 3.47 | 35.54 | 15 | 212.29 | 3 | 0 | 0 | 1 | 211.78 |
| Cyclohexanone, 3butyl | 3.16 | 17.07 | 11 | 154.25 | 1 | 0 | 0 | 3 | 171.78 |
| 2-Pentenoic acid, 3,4,4-trimethyl-, ethyl ester | 3.32 | 26.30 | 12 | 170.25 | 2 | 0 | 0 | 4 | 184.13 |
| 1,1'-Bicyclooctyl | 6.91 | 0.00 | 16 | 222.42 | 0 | 0 | 1 | 1 | 259.83 |
| 2H-Pyran-2carboxylic acid, 6-ethoxy-3,6-dihydro-, ethyl ester | 1.51 | 44.77 | 14 | 200.23 | 4 | 0 | 0 | 5 | 192.33 |
| Hexadecanoic acid, octyl ester | 9.34 | 26.30 | 26 | 368.65 | 2 | 0 | 1 | 22 | 426.56 |
| 2H-Pyran-2carboxylic acid, 6-butoxy-3,6-dihydro-, ethyl ester | 2.57 | 44.77 | 16 | 228.29 | 4 | 0 | 0 | 7 | 225.93 |
| 1,2-Cyclohexanedicarboxylic acid, didecyl ester | 9.42 | 52.61 | 32 | 452.72 | 4 | 0 | 1 | 22 | 494.15 |
| 6,10-Dimethyl-9undecen-4-ol | 4.58 | 20.23 | 14 | 198.35 | 1 | 1 | 0 | 7 | 231.98 |

Nearly half of the compounds violated Lipinski’s rule due to their higher molecular weights, reducing the oral drug usage. Yet, the compounds showed favourable properties for topical use because of their large volume and non-volatility. In addition, their low hydrogen bond donor and acceptor counts (nOHNH ≤1; nON ≤4) suggest moderate aqueous solubility and good membrane permeability. Especially, linoleic and oleic acid showed the lowest hydrogen bond donor (1) and acceptor (2) counts, with moderate TPSA, making them possess dermal permeability and enhancing interaction with bacterial membranes (Yuan. G, et al., 2021). Despite violating lipinski’s rule or having high molecular weight, these compounds are already FDA-approved excipients and extensively utilized in topical formulations.

#### 3.6.2. Pharmacokinetics analysis: (SwissADME)

The suitability of bioactive compounds, with key focus for topical application is further analysed on assessing pharmacokinetic properties by prioritizing low systemic absorption, not a BBB permeant, minimal cytochrome P450 inhibition and efflux transporters interaction, and a favourable skin permeability (Raney. S. G, et al., 2015).

Two thirds of the compounds exhibited high gastrointestinal (GI) absorption, while larger esters like hexadecanoic acid, octyl ester and didecyl cyclohexanedicarboxylate showed low GI absorption, making them less favourable for oral usage. Linoleic Acid had BBB permeant nature and inhibited CYP2C9, however, their common usage and good LogKp value makes it applicable for topical related usage. Several compounds (cyclohexanone and pyran esters) were predicted to cross the blood brain barrier, indicating possible CNS activity and neuropathic wound management. On the other hand, only 1,2-Cyclohexanedicarboxylic acid, didecyl ester was identified as a P-glycoprotein (P-gp) substrate possibly reducing intracellular accumulation.

Compounds like palmitic acid and nonyl laurate inhibited CYP1A2 and CYP2C9, potentially increasing the risk of drug–drug interactions, while ligands like benzodioxin and pyran esters showed no inhibition, making them safer option. Hexadecanoic acid, octyl ester had the Log Kp value of –0.85 cm/s, making it highly permeable to skin and ideal for dermal use, while compounds having Log Kp < –5.0 cm/s indicates slower skin absorption (Raney. S. G, et al., 2015).

**Table.6.**
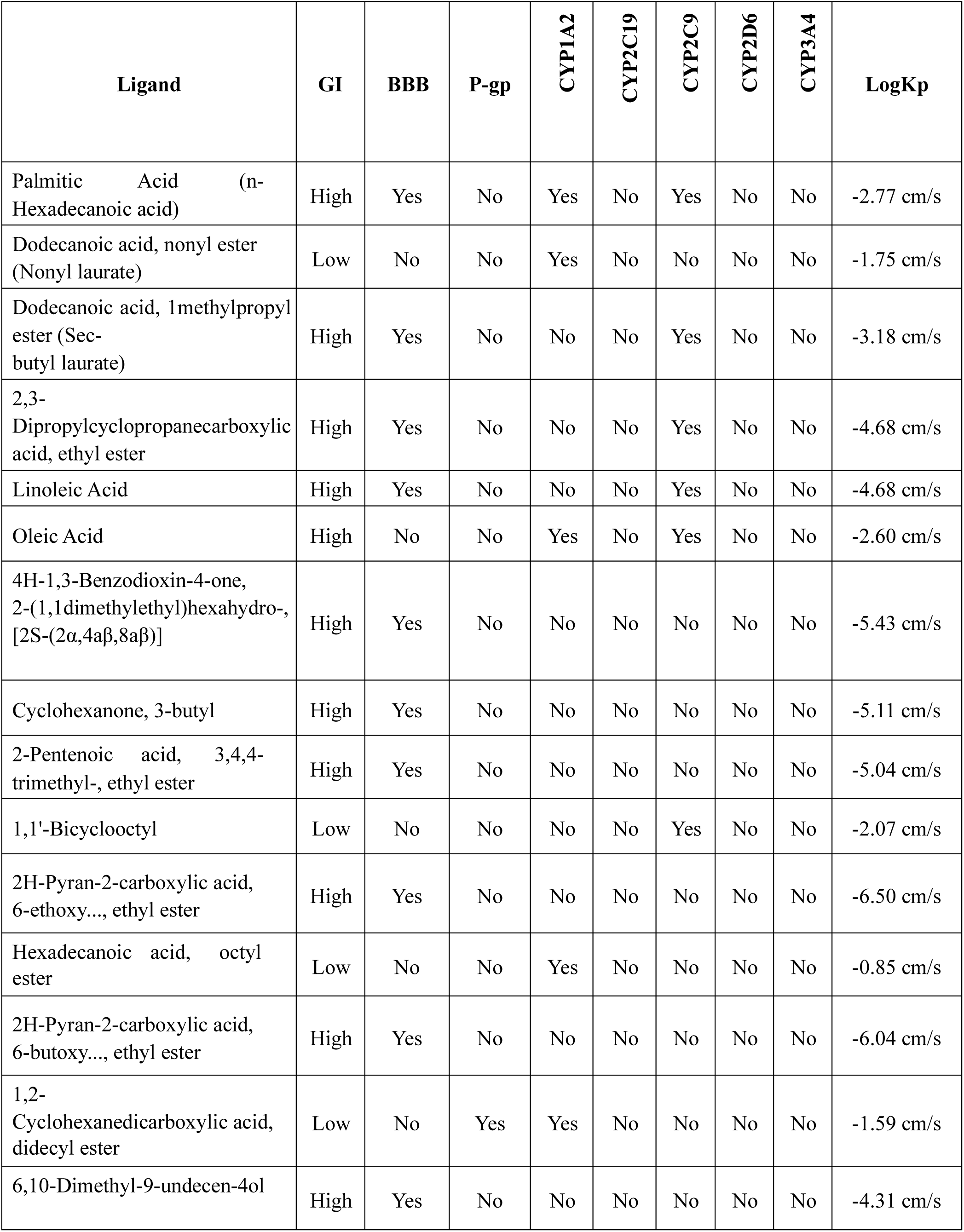
Drug-Likeness and Metabolic Interaction Analysis of Bioactive Compounds from Plant extract preserved at low temperature.

| Ligand | GI | BBB | P-gp | CYP1A2 | CYP2C19 | CYP2C9 | CYP2D6 | CYP3A4 | LogKp |
| --- | --- | --- | --- | --- | --- | --- | --- | --- | --- |
| Palmitic Acid (n-Hexadecanoic acid) | High | Yes | No | Yes | No | Yes | No | No | -2.77 cm/s |
| Dodecanoic acid, nonyl ester (Nonyl laurate) | Low | No | No | Yes | No | No | No | No | -1.75 cm/s |
| Dodecanoic acid, 1methylpropyl ester (Sec-butyl laurate) | High | Yes | No | No | No | Yes | No | No | -3.18 cm/s |
| 2,3-Dipropylcyclopropanecarboxylic acid, ethyl ester | High | Yes | No | No | No | Yes | No | No | -4.68 cm/s |
| Linoleic Acid | High | Yes | No | No | No | Yes | No | No | -4.68 cm/s |
| Oleic Acid | High | No | No | Yes | No | Yes | No | No | -2.60 cm/s |
| 4H-1,3-Benzodioxin-4-one, 2-(1,1dimethylethyl)hexahydro-, [2S-(2α,4$\alpha\beta$,8$\alpha\beta$)] | High | Yes | No | No | No | No | No | No | -5.43 cm/s |
| Cyclohexanone, 3-butyl | High | Yes | No | No | No | No | No | No | -5.11 cm/s |
| 2-Pentenoic acid, 3,4,4-trimethyl-, ethyl ester | High | Yes | No | No | No | No | No | No | -5.04 cm/s |
| 1,1'-Bicyclooctyl | Low | No | No | No | No | Yes | No | No | -2.07 cm/s |
| 2H-Pyran-2-carboxylic acid, 6-ethoxy..., ethyl ester | High | Yes | No | No | No | No | No | No | -6.50 cm/s |
| Hexadecanoic acid, octyl ester | Low | No | No | Yes | No | No | No | No | -0.85 cm/s |
| 2H-Pyran-2-carboxylic acid, 6-butoxy..., ethyl ester | High | Yes | No | No | No | No | No | No | -6.04 cm/s |
| 1,2-Cyclohexanedicarboxylic acid, didecyl ester | Low | No | Yes | Yes | No | No | No | No | -1.59 cm/s |
| 6,10-Dimethyl-9-undecen-4ol | High | Yes | No | No | No | No | No | No | -4.31 cm/s |

#### 3.6.3. Toxicity Analysis

To assess the safety and applicability of compounds, LD₅₀ (lethal dose), toxicity class, and predicted carcinogenicity, immunotoxicity, mutagenicity, cytotoxicity, and clinical toxicity were analyzed (Banerjee. P, & Ulker. O. C., 2022). All bioactive compounds indicated predominantly nontoxic nature, making them safe for application in pharmaceutical and cosmeceutical formulations.

The predicted LD₅₀ values ranged from 48 mg/kg to 60,000 mg/kg, with toxicity class present from 4 to 6 for all compounds except oleic acid, which is under toxicity class 2 sue to its lowest LD₅₀ (48 mg/kg). On the other hand, 1,2-cyclohexanedicarboxylic acid, didecyl ester showed the highest LD₅₀ (60,000 mg/kg) making it a class 6 compound and ensuring its application for both oral and skinbased application. All 15 compounds were inactive for hepatotoxicity, while, 1’-bicyclooctyl (0.57) and cyclohexanone, 3-butyl (0.50) showed possible neurotoxicity. Cardiotoxicity predictions were predicted to be negative for all compounds with high scores (≥0.70), indicating low risk of cardiac complications. Similarly, all compounds were predicted to be inactive for immunotoxicity, mutagenicity and cytotoxicity, with compounds like palmitic acid, oleic acid, linoleic acid and sec-butyl laurate getting perfect scores (1.0) for non-mutagenicity. While the majority were non-carcinogenic, four compounds showed mild carcinogenicity (<0.75), suggesting caution when used for topical applications to minimize systemic exposure.

**Table.7.** In Silico Prediction of Organ-Specific Toxicities for Selected Ligands from Plant extract stored at ambient temperature.

| Compound | Predicted LD<sub>50</sub><br>mg/kg | Toxicity Class | Hepatotoxicity | Neurotoxicity | Cardiotoxicity | Carcinogenicity | Immunotoxicity | Mutagenicity | Cytotoxicity |
| --- | --- | --- | --- | --- | --- | --- | --- | --- | --- |
| Palmitic Acid<br>(nHexadecanoic acid) | 900 | 4 | Inactive<br>(0.52) | Inactive<br>(0.92) | Inactive<br>(0.99) | Inactive<br>(0.63) | Inactive<br>(0.99) | Inactive<br>(1.0) | Inactive<br>(0.74) |
| Dodecanoic<br>Acid, Nonyl Ester (Nonyl<br>Laurate) | 5000 | 5 | Inactive<br>(0.76) | Inactive<br>(0.87) | Inactive<br>(0.79) | Active<br>(0.56) | Inactive<br>(0.96) | Inactive<br>(0.99) | Inactive<br>(0.78) |
| Dodecanoic<br>Acid, 1-Methylpropyl<br>Ester (Sec-butyl<br>Laurate) | 5000 | 5 | Inactive<br>(0.57) | Inactive<br>(0.81) | Inactive<br>(0.86) | Inactive<br>(0.51) | Inactive<br>(0.99) | Inactive<br>(0.99) | Inactive<br>(0.65) |
| 2,3-Dipropylcyclopropanecar<br>boxylic acid,<br>ethyl ester | 5000 | 5 | Inactive<br>(0.69) | Inactive<br>(0.77) | Inactive<br>(0.53) | Active<br>(0.54) | Inactive<br>(0.99) | Inactive<br>(0.79) | Inactive<br>(0.76) |
| Linoleic Acid<br>(9,12-Octadecadienoic acid<br>(Z,Z)-) | 10000 | 6 | Inactive<br>(0.55) | Inactive<br>(0.91) | Inactive<br>(0.99) | Inactive<br>(0.64) | Inactive<br>(0.96) | Inactive<br>(1.0) | Inactive<br>(0.71) |
| Oleic Acid | 48 | 2 | Inactive<br>(0.55) | Inactive<br>(0.91) | Inactive<br>(0.99) | Inactive<br>(0.64) | Inactive<br>(0.99) | Inactive<br>(1.0) | Inactive<br>(0.71) |
| 4H-1,3-Benzodioxin-4-one,<br>2-(1,1-dimethylethyl)hexahydro- | 1600 | 4 | Inactive<br>(0.78) | Inactive<br>(0.72) | Inactive<br>(0.64) | Inactive<br>(0.54) | Inactive<br>(0.99) | Inactive<br>(0.72) | Inactive<br>(0.81) |
| Cyclohexanone,<br>3-butyl | 500 | 4 | Inactive<br>(0.74) | Active<br>(0.50) | Inactive<br>(0.71) | Inactive<br>(0.75) | Inactive<br>(0.97) | Inactive<br>(0.88) | Inactive<br>(0.66) |
| 2 Pentenoic acid, 3,4,4-trimethyl-, ethyl ester | 3000 | 5 | Inactive<br>(0.73) | Inactive<br>(0.73) | Inactive<br>(0.82) | Active<br>(0.51) | Inactive<br>(0.99) | Inactive<br>(0.89) | Inactive<br>(0.76) |
| 1,1'-Bicyclooctyl | 750 | 3 | Inactive<br>(0.82) | Active<br>(0.57) | Inactive<br>(0.81) | Inactive<br>(0.50) | Inactive<br>(0.99) | Inactive<br>(0.92) | Inactive<br>(0.80) |
| 2H-Pyran-2carboxylic acid,<br>6-ethoxy-3,6dihydro-, ethyl ester | 590 | 4 | Inactive<br>(0.79) | Inactive<br>(0.82) | Inactive<br>(0.57) | Inactive<br>(0.60) | Inactive<br>(0.97) | Inactive<br>(0.67) | Inactive<br>(0.78) |
| Hexadecanoic acid, octyl ester | 5000 | 5 | Inactive<br>(0.76) | Inactive<br>(0.87) | Inactive<br>(0.79) | Active<br>(0.56) | Inactive<br>(0.96) | Inactive<br>(0.99) | Inactive<br>(0.78) |
| 2H-Pyran-2carboxylic acid,<br>6-butoxy-3,6dihydro-, ethyl ester | 590 | 4 | Inactive<br>(0.78) | Inactive<br>(0.82) | Inactive<br>(0.57) | Inactive<br>(0.55) | Inactive<br>(0.86) | Inactive<br>(0.73) | Inactive<br>(0.80) |
| 1,2-Cyclohexanedicarboxylic acid,<br>didecyl ester | 60000 | 6 | Inactive<br>(0.84) | Inactive<br>(0.85) | Inactive<br>(0.73) | Active<br>(0.72) | Inactive<br>(0.99) | Inactive<br>(0.98) | Inactive<br>(0.89) |
| 6,10-Dimethyl-9-undecen-4-ol | 5000 | 5 | Inactive (0.74) | Inactive (0.75) | Inactive (0.91) | Inactive (0.68) | Inactive (0.86) | Inactive (0.94) | Inactive (0.80) |

Acute toxicity data using the GUSAR model for the bioactive compounds showed favourable safety profile for topical or localized subcutaneous applications. 13 out of 15 compounds were predicted to be either non-toxic or OECD Class 5 and within the applicability domain (In AD). This indicates reliable prediction of low acute toxicity for both oral and SC routes. Compounds Nonyl laurate, Hexadecanoic acid, octyl ester and Sec-butyl laurate have very high SC LD₅₀ values of 15240, 17170 and 10360 mg/kg respectively, indicating exceptional safety for subcutaneous route. Likewise, essential fatty acids like palmitic acid, oleic acid and linoleic acid showed SC LD₅₀ values >3500 mg/kg, while compounds 2,3-dipropyl-cyclopropanecarboxylic acid, benzodioxin derivative, cyclohexanone and 1,1’-bicyclooctyl were classified under OECD Class 4 due to low LD_50_ values, making them moderately toxic (Ghannay. S, et al., 2020; Utaganovich. F. S, & Ismatovich. B. K., 2023).

For oral administration, most compounds were predicted to have very low toxicity, especially compounds like 1,2-cyclohexanedicarboxylic acid, didecyl ester and nonyl laurate had 10110 mg/kg and 8195 mg/kg LD_50_ values respectively, supporting the possibility for nutraceutical and therapeutic applications. Compounds with Class 5 ratings for both routes (e.g., 2H-pyran derivatives, 2-pentenoic acid ester) show promise for balanced systemic and topical application, controlled-release and feasibility, thereby validating them as preclinically safe.

**Table.8.** Assessment of Subcutaneous Route-Specific Acute Toxicity and Hazard Classification of Select Organic Molecules from Plant extract stored at ambient temperature.

| Compound | Route | Log<sub>10</sub> (LD<sub>50</sub>, mmol/kg) | LD<sub>50</sub> (mg/kg) | Applicability Domain (AD) | OECD Toxicity Classification |
| --- | --- | --- | --- | --- | --- |
| **Palmitic Acid (n-Hexadecanoic acid)** | Oral | 1.194 | 4010 | In AD | Class 5 |
|  | SC | 1.225 | 4306 | In AD | Non-Toxic |
| **Nonyl Laurate (Dodecanoic acid, nonyl ester)** | Oral | 1.4 | 8195 | In AD | Non-Toxic |
|  | SC | 1.669 | 15240 | In AD | Non-Toxic |
| **Sec-butyl Laurate (Dodecanoic acid, 1methylpropyl ester)** | Oral | 1.343 | 5647 | In AD | Non-Toxic |
|  | SC | 1.606 | 10360 | In AD | Non-Toxic |
| **2,3-Dipropylcyclopropanecarboxylic acid, ethyl ester** | Oral | 1.37 | 4653 | In AD | Class 5 |
|  | SC | 0.7 | 992.9 | In AD | Class 4 |
| **Linoleic Acid (9,12-Octadecadienoic acid (Z,Z)-)** | Oral | 1.273 | 5259 | In AD | Non-Toxic |
|  | SC | 1.376 | 6663 | In AD | Non-Toxic |
| **Oleic Acid** | Oral | 1.273 | 5302 | In AD | Non-Toxic |
|  | SC | 1.1 | 3555 | In AD | Non-Toxic |
| **4H-1,3-Benzodioxin-4one, 2-(1,1dimethylethyl)hexahydro-, [2S-(2α,4α $\beta$,8α $\beta$)]** | Oral | 0.947 | 1881 | In AD | Class 4 |
|  | SC | 0.228 | 358.7 | In AD | Class 4 |
| **Cyclohexanone, 3-butyl** | Oral | 1.251 | 2747 | In AD | Class 5 |
|  | SC | 0.546 | 542.8 | In AD | Class 4 |
| **2-Pentenoic acid, 3,4,4trimethyl-, ethyl ester** | Oral | 1.44 | 4689 | In AD | Class 5 |
|  | SC | 1.005 | 1724 | In AD | Class 5 |
| **1,1'-Bicyclooctyl** | Oral | 1.528 | 7509 | In AD | Non-Toxic |
|  | SC | 0.557 | 802.5 | In AD | Class 4 |
| **2H-Pyran-2-carboxylic acid, 6-ethoxy-3,6dihydro-, ethyl ester** | Oral | 1.236 | 3445 | In AD | Class 5 |
|  | SC | 0.994 | 1977 | In AD | Class 5 |
| **Hexadecanoic acid, octyl ester** | Oral | 1.37 | 8643 | In AD | Non-Toxic |
|  | SC | 1.668 | 17170 | In AD | Non-Toxic |
| **2H-Pyran-2-carboxylic acid, 6-butoxy-3,6dihydro-, ethyl ester** | Oral | 1.103 | 2894 | In AD | Class 5 |
|  | SC | 0.903 | 1827 | In AD | Class 5 |
| **1,2-Cyclohexanedicarboxylic acid, didecyl ester** | Oral | 1.349 | 10,110.00 | In AD | Non-Toxic |
|  | SC | 1.243 | 7915 | In AD | Non-Toxic |
| **6,10-Dimethyl-9-undecen-4-ol** | Oral | 1.429 | 5332 | In AD | Non-Toxic |
|  | SC | 0.732 | 1069 | In AD | Class 5 |

In addition to subcutaneous toxicity, the safety profiles of the compounds were comparatively analyzed using VEGA to confirm their mutagenic potential, skin compatibility, biodegradability and systemic risk. Except 2H-pyran-2-carboxylic acid (6-ethoxy derivative), every other compound was predicted to be non-mutagenic with scores ≥0.75, reflecting a low genotoxic risk. Cyclohexanone and pyran compounds showed mild irritation potential suggesting caution when incorporated into formulations. Rest of the compounds were classified as potential sensitizers, particularly fatty acids and esters (palmitic acid, nonyl laurate, and sec-butyl laurate), though skin irritation was minimum. Compounds such as 1,2-cyclohexanedicarboxylic acid, didecyl ester, and hexadecanoic acid octyl ester show excellent dermal tolerance, are non-sensitizers and non-irritants (Gaynanova. G, et al., 2021).

Majority of compounds were predicted to be non-irritating to ocular tissues and the prediction is supported by both experimental (EXP) and model basis. Linoleic acid showed eye irritation (GR) in one model, making it suitable for cutaneous use only. Biodegradability was observed for 80% of the compounds as they are readily biodegradable and skin permeation values (LogKp) ranging between – 2.9 cm/s to 1.99 cm/s suggests low systemic absorption of bioactive compounds, a desirable trait for topical wound agents. Benzodioxin, pyran esters and pentenoic acids have negative LogKp values indicating their low percutaneous absorption, while, fatty acid esters showed higher LogKp, indicating better skin penetration and suitability for deep tissue wound formulations (Gaynanova. G, et al., 2021). Total body elimination half-lives support their safe excretion profile and the data suggests that palmitic acid, hexadecanoic acid octyl ester, didecyl ester, oleic and linoleic acid to be highly suitable compounds for topical application (Maculewicz. J, et al., 2022).

**Table.9.** Toxicological and Dermal Safety Profiling (Predictability: LR - Low Reliability of Prediction, MR – Moderate Reliability of Prediction and GR – Good Reliability of Prediction, EXP - Experimental Value) of DCM-Extracted Compounds for Topical Application.

| Ligands | Mutag<br>enicity<br>(Ames<br>Test) | Skin Sensitization |  |  | Skin Irritation |  |  | Eye<br>Irrit<br>ation | Ready<br>Biodegr<br>adabilit<br>y | Skin<br>Perm<br>eatio<br>n<br>(Log<br>Kp) | Total<br>Body<br>Elimi<br>natio<br>n |
| --- | --- | --- | --- | --- | --- | --- | --- | --- | --- | --- | --- |
|  |  |  |  |  |  |  |  |  |  |  | Halfli<br>f |
|  | CONSENSUS | CAESAR | TOXTREE | CONCERT/Kode | CONCERT/Kode | CONCERT/SarPy | EXPERIMENTAL | CONCERT/Kode | IRFMN | Potts and Guy | QSARINS |
| Palmitic Acid<br>(or)<br>n-Hexadecanoic acid | NON-Mutag<br>enic<br>(Cons<br>ensus<br>score:<br>1) | Sen<br>s<br>itize<br>r<br>(GR<br>) | No<br>clas<br>s<br>foun<br>d | Ina<br>ctiv<br>e<br>(MR) | NO<br>N-Sen<br>s<br>itize<br>r<br>(MR<br>) | Irri<br>t<br>atin<br>g<br>(MR) | NOT<br>IRRI<br>TAN<br>T | NO<br>N-Irrita<br>ting<br>(EX<br>P) | Possible<br>Readily<br>Biodegr<br>a dable<br>(GR) | 0.82<br>7<br>(LR) | 0.449<br>(MR) |
| Dodecanoic acid,<br>nonyl ester<br>(or)<br>Nonyl laurate | NON-Mutagenic<br>(Consensus score: 0.75) | Sensitizer<br>(GR) | No class found | Active<br>(GR) | NON-Sensitizer<br>(GR) | Irritating<br>(LR) | NOT IRRITANT<br>(GR) | NON-Irritating<br>(GR) | Readily Biodegradable<br>(MR) | 1.275<br>(LR) | 0.649<br>(MR) |
| Dodecanoic acid, 1-methylpropyl ester<br>(or)<br>Sec-butyl laurate | NON-Mutagenic<br>(Consensus score: 0.9) | Sensitizer<br>(GR) | No class found | Active<br>(GR) | NON-Sensitizer<br>(MR) | Irritating<br>(MR) | NOT IRRITANT<br>(GR) | NON-Irritating<br>(GR) | Readily Biodegradable<br>(GR) | 0.027<br>(LR) | 0.715<br>(MR) |
| 2,3-Dipropylcyclopropane carboxylic acid,<br>ethyl ester | NON-Mutagenic<br>(Consensus score: 0.825) | Sensitizer<br>(GR) | No class found | Active<br>(MR) | NON-Sensitizer<br>(LR) | NON-Irritating<br>(LR) | NOT IRRITANT<br>(GR) | NON-Irritating<br>(GR) | NON Readily Biodegradable<br>(MR) | - 1.455<br>(MR) | 0.623<br>(MR) |
| 9,12-Octadecadienoic acid<br>(Z,Z)-<br>(or)<br>Linoleic Acid | NON-Mutagenic<br>(Consensus score: 1) | Sensitizer<br>(GR) | No class found | Active<br>(EXP) | NON-Sensitizer<br>(MR) | Irritating<br>(LR) | NOT IRRITANT<br>(MR) | Irritating<br>(GR) | Possible Readily Biodegradable<br>(MR) | 0.595<br>(LR) | 0.501<br>(LR) |
| Oleic Acid | NON-Mutagenic<br>(Consensus score: 1) | Sensitizer<br>(GR) | No class found | Active<br>(EXP) | NON-Sensitizer<br>(MR) | Irritating<br>(MR) | NOT IRRITANT<br>(MR) | NON-Irritating<br>(MR) | Possible Readily Biodegradable<br>(MR) | 1.002<br>(LR) | 0.473<br>(MR) |
| 4H-1,3Benzodioxin-4-one, 2-(1,1dimethylethyl)hexahydro-, [2S-(2.alpha.,4a.beta.,8a.beta.)] | NON-Mutagenic (Consensus score: 0.575) | Sensitizer (MR) | No class found | Inactive (LR) | NO N-Sensitizer (LR) | NO N-Irritating (LR) | NOT IRRITANT (GR) | N/A | Readily Biodegradable (LR) | - 1.909 (LR) | 0.759 (MR) |
| Cyclohexanone, 3-butyl | NON-Mutagenic (Consensus score: 0.575) | Sensitizer (GR) | No class found | Active (MR) | Sensitizer (MR) | Irritating (LR) | IRRITANT (LR) | NO N-Irritating (GR) | Possible Readily Biodegradable (MR) | - 1.656 (MR) | 0.627 (MR) |
| 2-Pentenoic acid, 3,4,4trimethyl-, ethyl ester | NON-Mutagenic (Consensus score: 0.9) | Sensitizer (GR) | Schiff Base Formation | Inactive (LR) | Sensitizer (MR) | Irritating (LR) | NOT IRRITANT (MR) | N/A | Possible Readily Biodegradable (MR) | - 1.729 (MR) | 0.987 (MR) |
| 1,1'-Bicyclooctyl | NON-Mutagenic (Consensus score: 0.825) | Sensitizer (GR) | No class found | Active (MR) | Sensitizer (LR) | Irritating (LR) | NOT IRRITANT (GR) | NO N-Irritating (GR) | NON Readily Biodegradable (LR) | 0.572 (LR) | 1.19 (MR) |
| 2H-Pyran-2carboxylic acid, 6-ethoxy-3,6dihydro-, ethyl ester | Mutagenic (Consensus score: 0.15) | Sensitizer (MR) | No class found | Active (LR) | NO N-Sensitizer (MR) | Irritating (LR) | IRRITANT (MR) | N/A | Readily Biodegradable (MR) | - 2.945 (MR) | 0.587 (MR) |
| Hexadecanoic acid, octyl ester | NON-Mutagenic (Consensus score: 0.75) | Sensitizer (GR) | No class found | Inactive (LR) | NO N-Sensitizer (GR) | Irritating (LR) | NOT IRRITANT (GR) | NO N-Irritating (GR) | Readily Biodegradable (GR) | 1.99 (LR) | 0.649 (MR) |
| 2H-Pyran-2carboxylic acid, 6-butoxy-3,6dihydro-, ethyl ester | NON-Mutagenic (Consensus score: 0.25) | Sensitizer (MR) | No class found | Active (MR) | Sensitizer (MR) | Irritating (LR) | NOT IRRITANT (MR) | N/A | Readily Biodegradable (MR) | - 2.421 (LR) | 0.647 (MR) |
| 1,2-Cyclohexanedicarboxylic acid, didecyl ester | NON-Mutagenic (Consensus score: 0.675) | Sensitizer (GR) | No class found | Inactive (MR) | NO N-Sensitizer (GR) | NO N-Irritating (GR) | NOT IRRITANT (GR) | NO N-Irritating (GR) | NON Readily Biodegradable (MR) | 1.419 (LR) | 0.586 (MR) |
| 6,10-Dimethyl-9undecen-4-ol | NON-Mutagenic (Consensus score: 0.825) | Sensitizer (GR) | No class found | Active (GR) | NO N-Sensitizer (LR) | Irritating (LR) | NOT IRRITANT (MR) | NO N-Irritating (GR) | Possible Readily Biodegradable (GR) | - 0.781 (MR) | 0.803 (MR) |

### 3.7. Molecular Docking and Interaction Studies

#### 3.7.1. Prediction of Active Binding Site

Molecular docking was performed for polyisoprenyl-teichoic acid--peptidoglycan teichoic acid transferase protein to assess the binding affinity and interaction profiles of standard topical drugs. The Docking simulations were carried out by placing the Grid Center at x = -0.4527, y = -0.562, z = -0.8638, with Grid Size being x = 75.76, y = 122.14, z = 66.95, and the results are presented in Figure 7 and Table 10.

**Fig.7.**
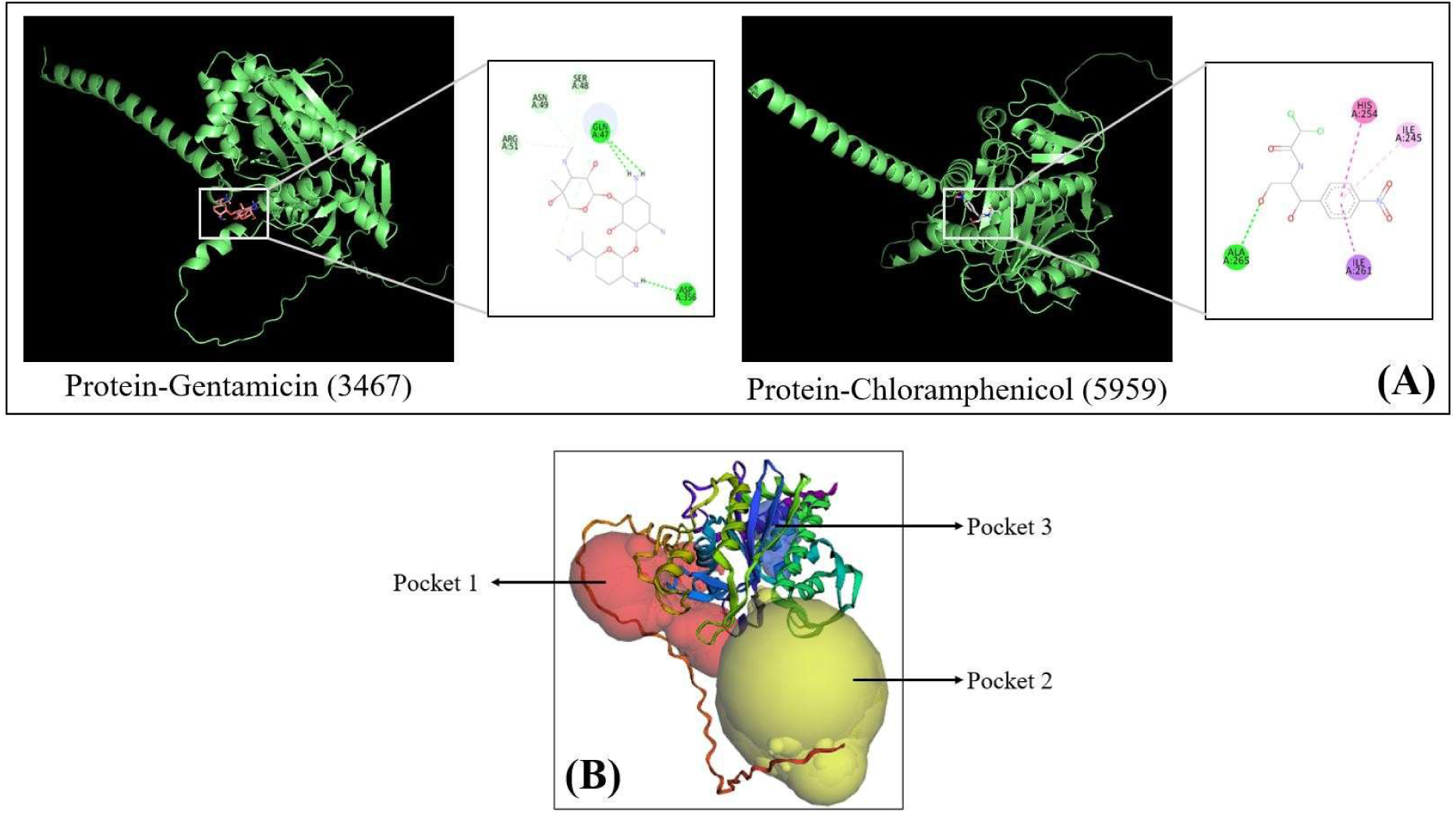
(A) Comparative Docking Analysis of Standard Drug Compounds with PTA and their Binding Interactions; (B) 3D Protein structure presenting the Predicted Binding Pockets on the Target Protein Surface using CastP.

**Table.10.** (a) Binding Affinities and Interaction Profiles of Drug Candidates from Algal Extracts and Standard Drugs with PTA; (b) Best Predicted Binding Pockets based on Surface Area, Volume, and Key Amino Acid Residues.

| **(a) Molecular Docking – Blind Docking using Standard Drugs** |  |  |  |  |
| --- | --- | --- | --- | --- |
| **Ligand** | **Binding Affinity** | **Interacting Protein Residues** | **No. of Bonds** | **Interaction Type** |
| **Chloramphenicol (5959)** | -6.5 | Ala(A)265 | 1 | Conventional Hydrogen Bonds |
|  |  | His(A)254 | 1 | Pi-Pi T-shaped |
|  |  | Ile(A)261 | 1 | Pi-Sigma |
|  |  | Ile(A)245 | 1 | Pi-Alkyl |
| Gentamicin (3467) | -6 | Gln(a)47<br>Asp(A)356 | 3 | Conventional Hydrogen Bonds |
|  |  | Ser(A)48<br>Asn(A)49<br>Arg(A)51, | 3 | Carbon-Hydrogen bonds |
| (b) CastP Pocket prediction |  |  |  |  |
| Pocket ID | Surface Area (Å²) | Volume (Å³) | Residue Count | Key Amino Acids in Binding Pocket |
| 1 | 1227.775 | 3776.364 | 52 | ASP40, LYS42, ASN43, GLY79, GLN80, AS81, GLU83, ARG86, TYR114, PRO116, LYS117, VAL118, GLY119, TYR120, HIS126, HIS128, ALA129, TYR130, GLY131, GLY132, PRO133, ILE134, TYR151, ARG153, LYS253, LEU337, PRO339, LEU340, SER342, LEU345, ARG346, GLN349, ASN350, ASP351, THR353, ASP354, ASP356, LYS357, SER358, ASN361, ASN372, SER373, VAL374, ASN375, ASN376, GLN377, GLN378, GLN379, PRO380, ALA381, THR382, ASP383 |
| 2 | 263.933 | 2278.219 | 24 | PRO108, ARG109, ASP110, ARG209, HIS210, HIS211, ASP212, SER213, ASP214, LEU215, ARG217, ASP284, MET286, TYR291, TYR293, SER400, ASN401, ASN402, GLN403, ASN404, GLY405, VAL406, VAL407, ASN408 |
| 3 | 522.417 | 407.678 | 39 | PHE35, ILE64, PHE66, LEU67, GLY68, ALA89, ILE91, PHE95, ILE102, MET104, VAL152, VAL154, MET156, ALA158, PHE159, ALA162, LEU166, LEU203, ALA206, GLN221, LEU224, ILE225, LEU228, PHE229, ALA232, LEU241, VAL244, ILE245, VAL248, GLY249, ALA252, HIS254, LEU256, ILE261, LEU264, ALA265, TYR268, LEU269, ILE275 |

Chloramphenicol showed a binding affinity of -6.5 kcal/mol, and was involved in key interactions with residues Ala265, His254, Ile261 and Ile245. These residues are within the conserved Cps2A domain (residues 62–310), indicating that interactions here could interfere directly with the functional core of the protein, thereby possessing the potential for disrupting its role in anchoring teichoic acids to the peptidoglycan layer (Selvaraj. C, et al., 2022). Gentamicin showed a slightly lower binding affinity (–6.0 kcal/mol) and bound to 5 residues present outside the Cps2A domain of the protein, suggesting that it might not directly interfere with enzymatic function but could still interfere with the protein structure and its interactions.

CASTp analysis of the target protein was performed to identify potential active binding sites. Among the multiple pockets predicted by CastP, the First (best) three binding pockets which had comparatively larger surface area and volume were chosen, since they possess higher potential for ligand accommodation and interaction (Tibaut. T, et al., 2016). Residues which were identified to be interacting with PTA through blind docking were compared against residues predicted by CastP. Pocket 1 showed the largest surface area and residue count, while Pockets 2 and 3 were smaller, yet also contained critical residues. This suggests the presence of key active or allosteric site. Pocket 3 may support lipophilic interactions (chloramphenicol binding), while Pocket 1 accommodates polar and charged residues (gentamicin binding), underscoring a dual target nature of the protein. A total of five amino acid residues (ALA265, HIS254, ILE261, ILE245 and ASP356) were found to be common in both docking and CastP data, and their presence within the conserved catalytic domain of Cps2A suggests their role in maintaining the potential function.

Active site residues were selected in PyRx and the grid box was defined (centered at x = 4.7496, y = -11.7643, z = 5.0159 with dimensions x = 30.5099, y = 9.7284, z = 24.9810). The results are summarized in Table 11 and the interactions are presented in Figure 8.

**Fig.8.**
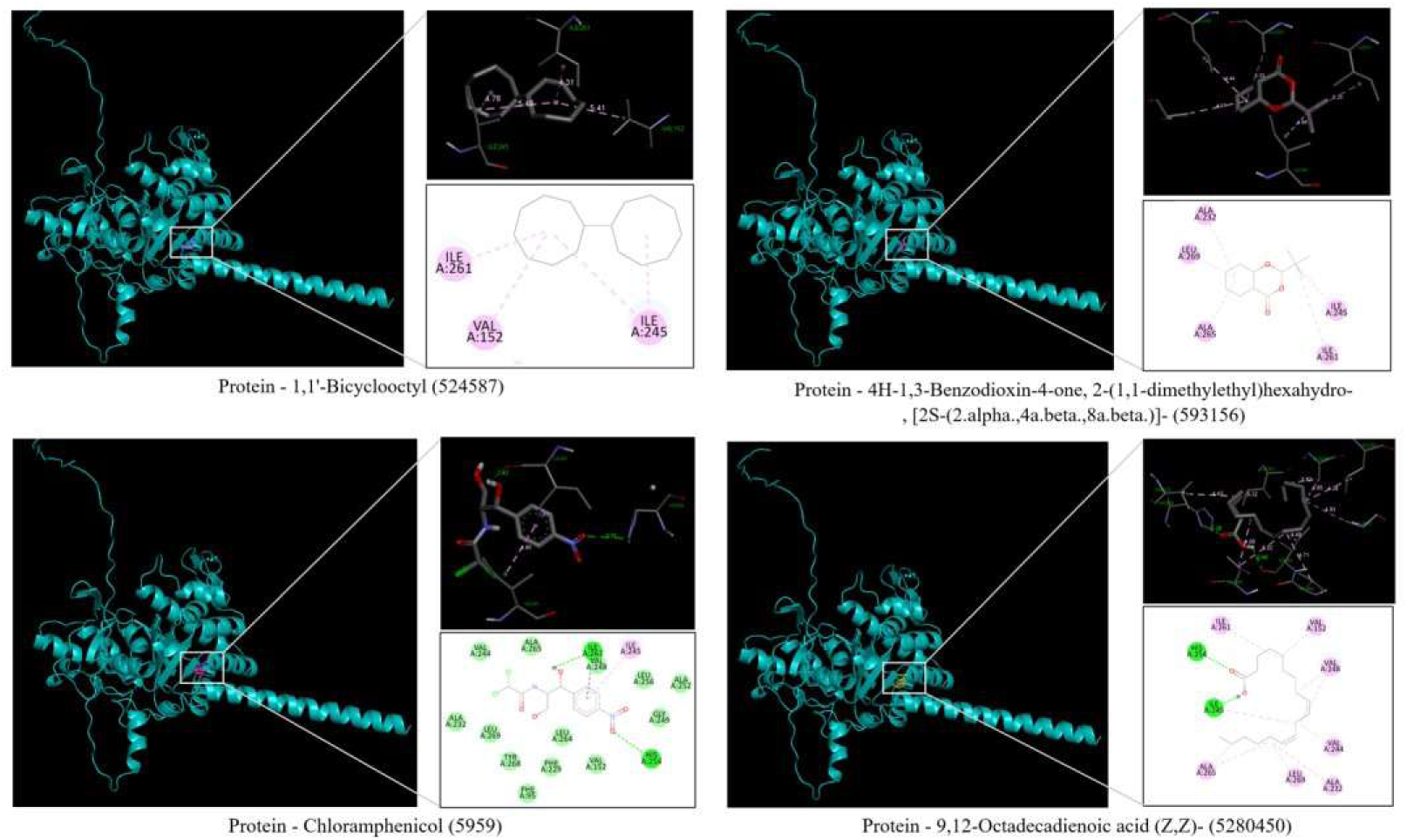
Active site Docking Analysis and Binding Interactions of Standard Drugs and Algal Bioactive Compounds with PTA.

**Table.11.**
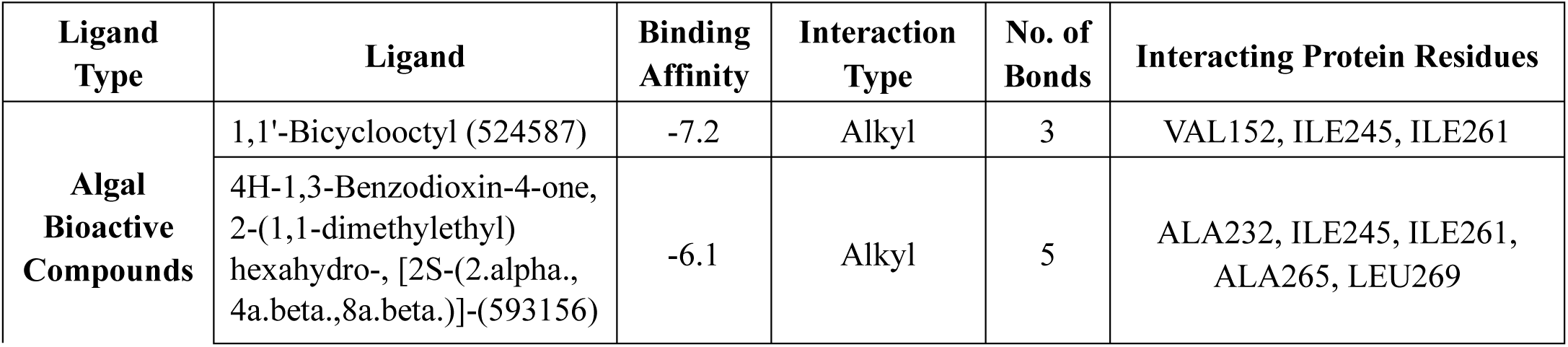
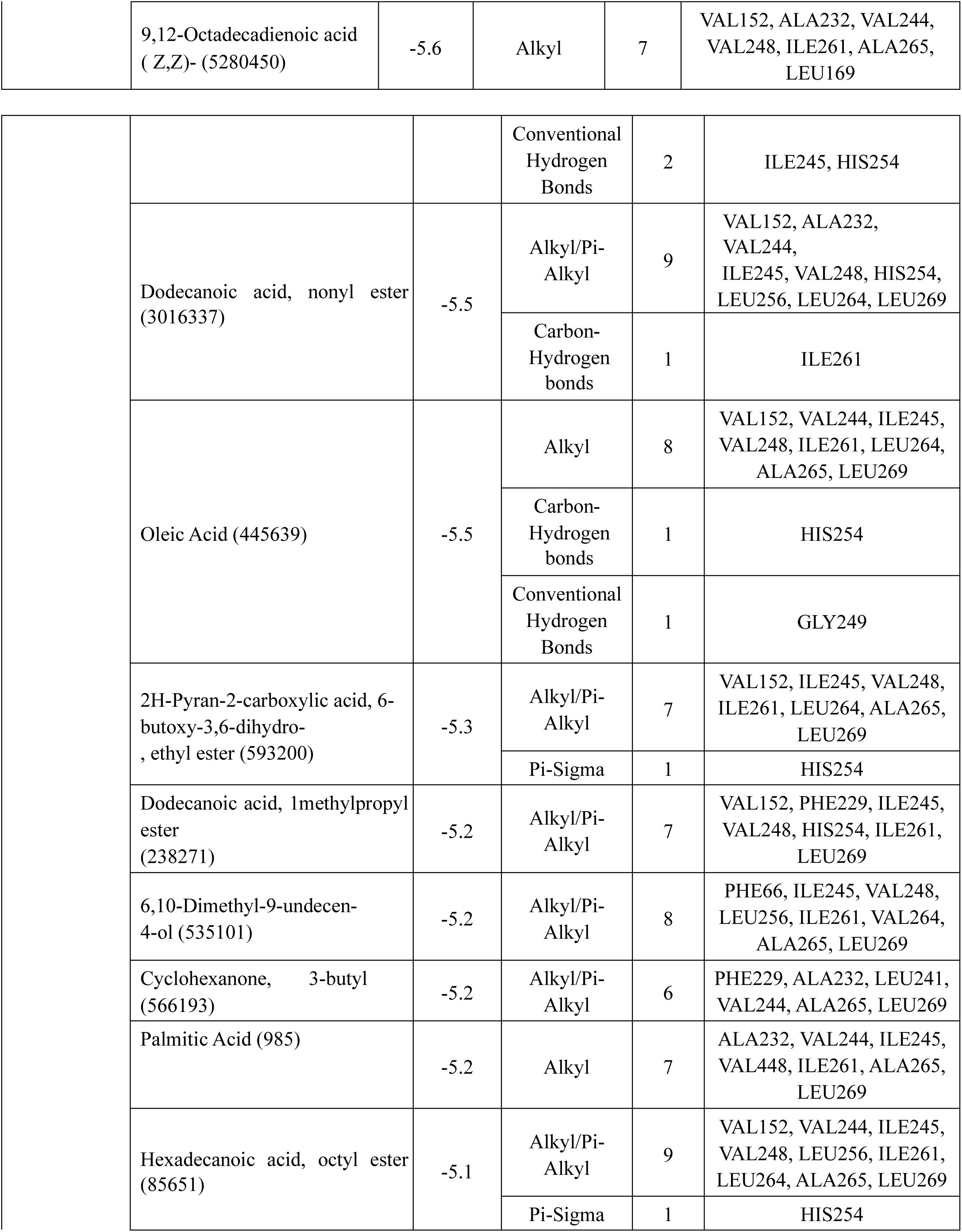
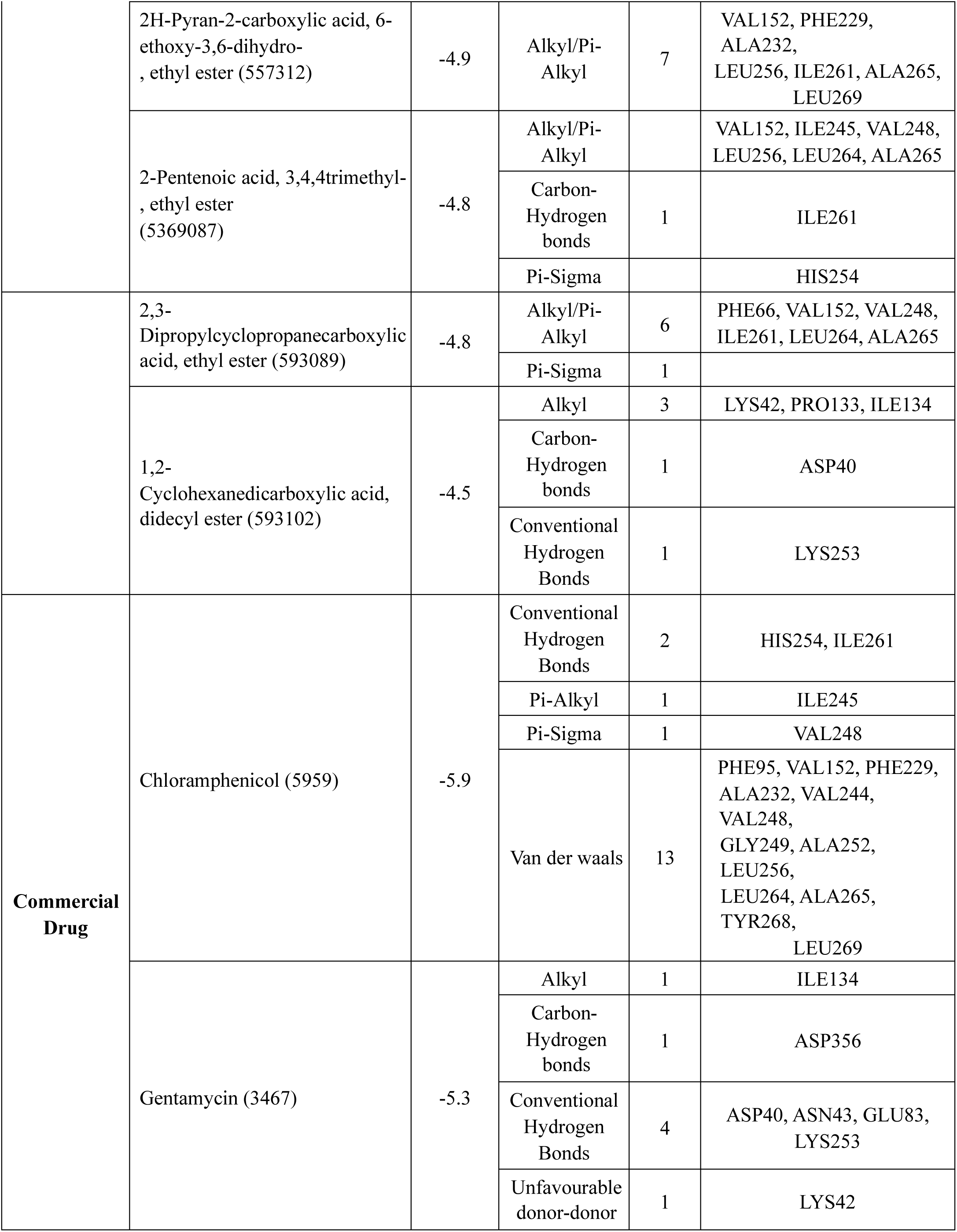
Binding Affinities and Interaction Profiles predicted using Active site Docking of Algal compounds and Standard Drugs.

The docking results revealed compound 1,1’-Bicyclooctyl (PubChem ID: 524587) to exhibit the highest binding affinity with a docking score of –7.2 kcal/mol. The interactions are primarily stabilized through alkyl interactions with key hydrophobic residues such as VAL152, ILE245, and ILE261, making the compound a promising drug candidate. Next to this, compound 4H-1,3Benzodioxin-4-one derivative (ID: 593156) demonstrated a binding affinity of –6.1 kcal/mol, with 5 alkyl interactions to residues ALA232, LEU269, etc., which are located near the active site. These two compounds were the only ones with a higher binding affinity than the control drug Chloramphenicol (– 5.9 kcal/mol) and gentamycin (–5.3 kcal/mol), suggesting that the ligands may engage effectively in inhibiting the protein’s biological function. Chloramphenicol in particular formed a diverse interactions like hydrogen bonds (HIS254, ILE261), π-alkyl and van der Waals interactions with VAL248, GLY249, TYR268 and multiple other residues, serving as the benchmark for comparison.

9,12-Octadecadienoic acid (Z,Z)-, i.e., linoleic acid showed a moderate affinity of –5.6 kcal/mol, but had 7 alkyl interactions and formed 2 conventional hydrogen bonds with ILE245 and HIS254. Dodecanoic acid, nonyl ester and Oleic acid both had –5.5 kcal/mol binding affinity, however, dodecanoic acid showed multiple alkyl and π-alkyl interactions with residues LEU256, LEU264 and HIS254, while oleic acid showed hydrogen bonding and hydrophobic interactions with residues GLY249 and HIS254. These interactions suggest fatty acids such as linoleic acid contribute to both structural and functional binding by interacting deeply within the binding pocket and act as membrane disruptors.

Compounds like 2H-Pyran-2-carboxylic acid derivatives and Dodecanoic acid, 1-methylpropyl ester had binding affinities better than -5.0 kcal/mol and involve alkyl/π-alkyl interactions with common residues (ALA265, ILE261, and LEU269). Moreover, most of the algal compounds matched the binding affinities of these standard drugs and consistently interacted with the active site residues, making them novel drug candidates for wound-healing and antibacterial applications.

An important note when targeting *S. aureus* is that the MRSA strains generally develop resistance to known antibiotic targets by altering them or activating efflux pumps (Lee. A. S, et al., 2018). Screening of *H. pluvialis* compounds through docking and ADMET studies identified several promising candidates having potential to be employed for a dual-targeting strategy for both oral and topical routes. Linoleic Acid, 2H-Pyran-2-carboxylic acid, 6-butoxy…, ethyl ester, and 4H-1,3Benzodioxin-4-one derivative showed the most favorable binding affinities and interactions with 4 key active site residues (ILE245, HIS254, ILE261, ALA265) making them comparable to or exceeding the control drug chloramphenicol.

Linoleic Acid is found to be the best drug candidate due to its strong binding (–5.6 kcal/mol), high safety margin and is already considered ideal for both oral and dermal routes. 4H-1,3-Benzodioxin4-one derivative came second best, even though having excellent affinity (–6.1 kcal/mol), due to class 4 toxicity and lesser LD_50_ value. However, the combination of both can support strong binding, dualroute compatibility and better safety profile, thereby ensuring feasibility to treat localized infections or inflammatory conditions, effectively enhancing antimicrobial efficacy (Feng. D, et al., 2023). This strategy impairs S. aureus’s function and minimize the chance of developing resistance, thereby presenting a promising aspect for targeting multidrug-resistant pathogens (Jiao. F, et al., 2024).

## 4. Conclusion

This study provides in-vitro analysis followed by a comprehensive in-silico characterization of polyisoprenyl-teichoic acid–peptidoglycan teichoic acid transferase (PTA), a key LCP family enzyme present in *Staphylococcus aureus* and essential for cell wall biosynthesis. The *H. pluvialis*-loaded gelpatch modified using acetone extract presented as safe and practical due to its antioxidant and antibacterial efficacy, and good stability. Despite a moderate reduction in antimicrobial diffusion of gel matrix compared to the extract, the patch’s safety (elimination of acetone via evaporation), customizability and biocompatibility make it a promising candidate for treating chronic S. aureus infected or oxidative-stress-related wounds.

In-silico validation of protein using domain analysis, physicochemical profiling, secondary and tertiary structure predictions confirm its structural aspects and potential to serve as a drug target potential, due to their absence in humans. GC-MS profiling of *H. pluvialis* extracts identified several bioactive compounds, with Linoleic Acid, 2H-Pyran-2-carboxylic acid, 6-butoxy…, ethyl ester and 4H1,3-Benzodioxin-4-one derivative being the lead candidates owing to its most favourable ADMET properties and strong binding affinities in the active site. These interactions with conserved residues present in the catalytic/active site of PTA suggest a potential inhibitory mechanism. In addition, utilizing this computational approach in identifying sustainable, topical antibacterial agents has found out the synergistic potential Linoleic Acid and 4H-1,3-Benzodioxin-4-one with standard antibiotics underscores a promising strategy against MRSA, and laying the groundwork for further experimental validation.

